# Sponges are highly resistant to radiation exposure and cancer

**DOI:** 10.1101/2021.03.17.435910

**Authors:** Angelo Fortunato, Jake Taylor, Jonathan Scirone, Athena Aktipis, Carlo C. Maley

## Abstract

There are no reports of cancer in sponges, despite them having somatic cell turnover, long lifespans and no specialized adaptive immune cells. In order to investigate whether sponges are cancer resistant, we exposed a species of sponge, *Tethya wilhelma*, to X-rays. We found that *T. wilhelma* can withstand 600 Gy of X-ray radiation. That is approximately 100 times the lethal dose for humans. A single high dose of X-rays did not induce cancer in sponges, providing the first experimental evidence of cancer resistance in the phylum, Porifera. Following X-ray exposure, we found an overexpression of genes involved in DNA repair, signaling transduction pathways and epithelial to mesenchymal transition. Sponges have the highest level of radiation resistance that has yet been observed in animals that have sustained somatic cell turnover. This may make them an excellent model system for studying cancer resistance and developing new approaches for cancer prevention and treatment.

## Introduction

To date there have been no reports of cancer in sponges^1^. Sponges are part of the phylum Porifera, and they have a long lifespan and somatic cell turnover^2^, which should make them susceptible to cancer because over the course of their lifespans they would be expected to accumulate carcinogenic mutations. Here we set out to investigate whether sponges are particularly cancer resistant, and, if so, what the mechanisms underlying this cancer resistance are. Through a combination of microscopy and transcriptomics, we were able to observe changes in *Tethya wilhelma* (Demosponges)^3,4^ after X-ray exposure and assess the organism-level, cell-level and gene expression changes over time.

Basal invertebrates like sponges that lack^5^ immune specialized cells^6^ or with primitive elements of an adaptive immune system^7,8^, may lack the ability to detect and eliminate mutant cells. They should be particularly susceptible to cancer. However, the fact that no cancer has been reported in sponges^1^ suggests that they might have a physiology that is resilient to mutations or possess effective mechanisms for DNA damage prevention, DNA repair, and tissue homeostasis.

### Sponges are an effective model system for studying cancer resistance

In order to investigate cancer resistance in sponges and evaluate this hypothesis, we studied the sponge *T. wilhelma*, which is a sessile, filter-feeding demosponge that originally came from the Indo-Pacific oceans^9^ (Fig.1). Demosponges possess a canal system which is characterized by a highly complex network of chambers lined with choanocytes, which are flagellated cells that are specialized in creating a flow of water and capturing food particles^10^. Water enters from the pores located mainly on the lateral walls of sponges and is discarded through a large medial excurrent canal opening on the apex of sponges, the osculum^10^. *T. wilhelma* have a globular shape and the largest specimens can reach the size of 15-20 mm in diameter (Fig.1). *T. wilhelma* reproduce asexually by budding in the laboratory^11^. These sponges are capable of contractile and slow locomotory behavior ^3,4^ and adapt well to being cultured in aquaria^3,4^.

**Figure 1.**
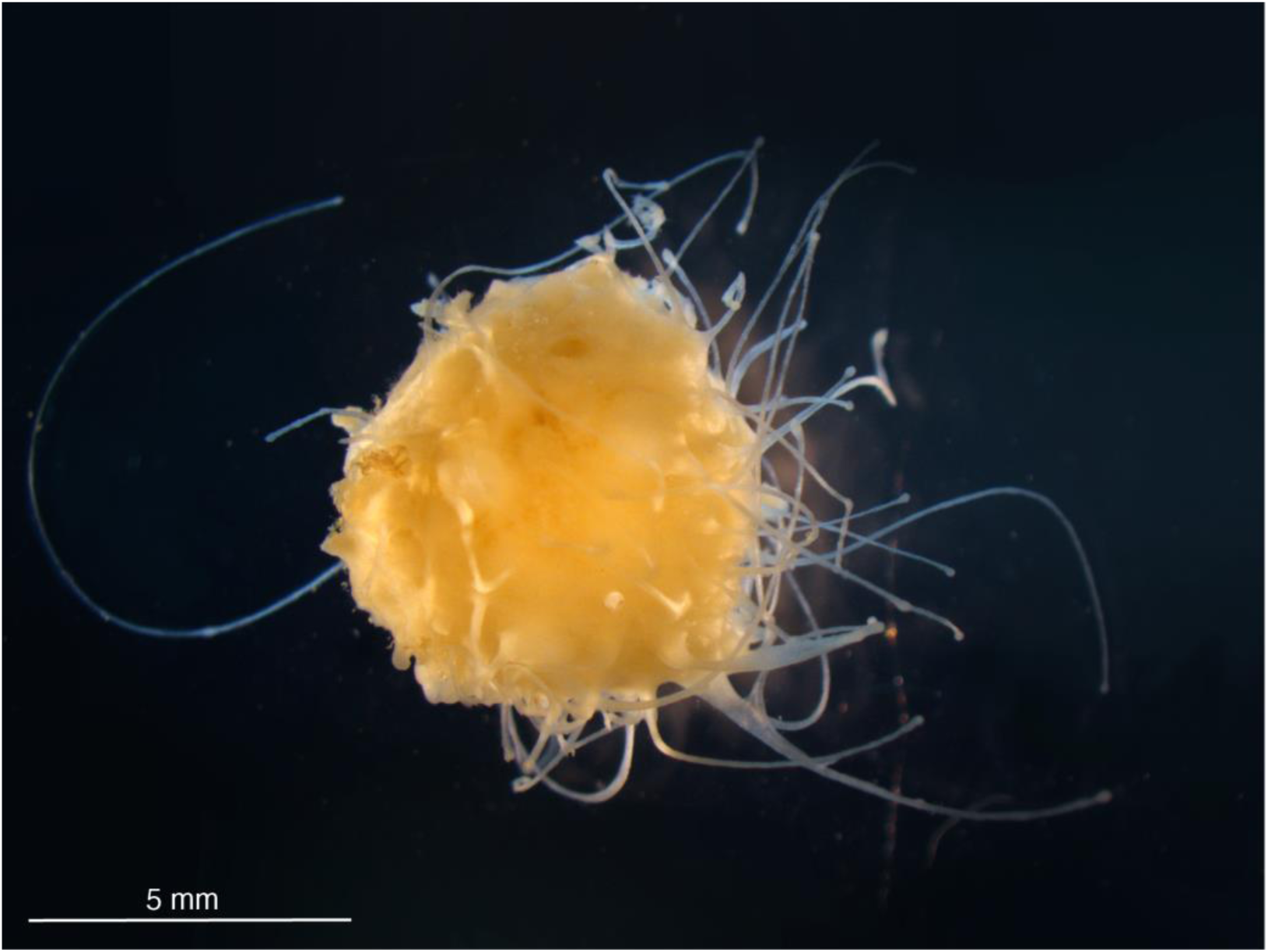
*T. wilhelma* have a globular shape. They produce filamentous body extensions and are not pigmented. The largest specimens can reach the size of 15-20 mm in diameter. This image was taken using dark field illumination.

The genome sequence of *T. wilhelma* is available^12^ and genomic analyses have shown that many molecular pathways (e.g. signal transduction mechanisms) are well conserved on sponges^12^. *T. wilhelma* has a remarkably long lifespan but sufficiently short generation time^3^ to make it possible to rapidly obtain experimental results.

This is the first study of radiation resistance and cancer resistance in porifera, the phylum at the very base of animal life on earth.

## Materials and Methods

### Lab cultures

We built a sponge culture system that consists of a main saltwater aquarium (340 liters) that develops an ecosystem (coral reef) capable of supporting the growth of the sponges. The sponges are grown in 3 smaller (30 liter) culture aquariums connected to the main one, with controlled temperature (24°C) and water flow. This setting allows easy access to the sponges, including the observation of the sponges under a microscope without having to remove them from the aquarium.

We fed the sponges with artificial plankton (Aquakultur Genzel GmbH, Germany) twice daily. In order to obtain fine (<25μm) food particles assimilable by sponges, we homogenized the artificial plankton with an IKA T10 Basic Ultra Turrax homogenizer.

The culturing of the sponges started 4 years prior to this study, with 20 animals. The original specimens are still alive, thus the lifespan of the sponges in our laboratory setting is > 4 years.

### DNA damage by X-ray irradiation

We exposed young adult sponges (diameter>5mm) to X-rays utilizing a RS-2000 Biological System X-ray irradiator. We exposed the sponges to a single dose of 600 Gy for the experiments. Considering X-ray absorbance of the 10 mm column of water above the animals during the X-ray irradiation, we estimated the actual X-ray exposure of specimens to be 13.7% lower, 518 Gy.

### Experimental settings

Based on preliminary morphological observations, we selected three time points: 24 hours, 7 days, and 21 days after X-ray exposure for both histological and transcriptomic analyses (RNA-seq). We randomly selected 3 sponges for each time point plus 3 specimens of control for both histological and transcriptome analyses, for a total of 36 sponges. We treated 4 additional independent groups of sponges (n=6, n=5, n=5, n=5, total 21 sponges) and relative controls (5 sponges for each group, total 20 sponges), for long-term morphological observations. Each group was exposed to X-rays at different times.

### Morphological analysis

We observed the morphological changes in the animals *in vivo* using ImageJ software^13^. Sponges are partially translucent, but only superficial structures can be observed *in vivo*. However, their shape is regular, and any morphological changes are easily observable.

For histological examination we fixed the specimens with Pampl’s fluid^14^ for 24 hours at 4°C. Then, we dissolved the siliceous spicules that make up its skeleton by submerging the specimens in 4% hydrofluoric acid (MilliporeSigma, cat. n. 1.00338) for an additional 24 hours at 4°C, then we followed standard histological protocols^15,16^.

For transmission electron microscopy, we fixed specimens in 2.5% glutaraldehyde (Electron Microscopy Sciences, cat.n. 16020) in 0.2 M Na-cacodylate sucrose buffer (pH 7.2; Electron Microscopy Sciences, cat.n. 12300) for 2.5 hours at 4°C. Then, we rinsed the specimens 3 times with a 0.2 cacodylate sucrose buffer for 45-60 min total, post fixed them for 2 hours in 1% osmium tetroxide (Electron Microscopy Sciences, cat n. 19150) 0.2 cacodylate sucrose buffer and washed them 1 time with buffer, then 3 times with deionized water for 45-60 min total. We stained them *en bloc* with 1% aqueous uranyl acetate (Electron Microscopy Sciences, cat n. 22400) for 16 hours at 4°C. After washing the specimens 4 times with water for 45-60 min total. We dehydrated them with an ascending ethanol series up to 70% ethanol. Then, we disilicate the specimens with 4% hydrofluoric acid for 1 hour at 4°C. Afterwards, we washed specimens in 70% ethanol, and we completed the dehydration with an ethanol series up to 100%. Then we transferred the specimens to anhydrous propylene oxide (cat. n. 14300) for 30 minutes (replacing the anhydrous propylene oxide with fresh one after 15 minutes). We infiltrated samples with 5% Spurr’s epoxy resin (in anhydrous propylene oxide 3 hours with rotation; 50% resin in anhydrous propylene oxide overnight with rotation (18 hr); 75% resin in anhydrous propylene oxide with rotation (6 hr); 100% pure resin 3x for 24 hr total (6 hr, 12 hr, 6 hr). Finally, we flat-embedded the specimens and polymerized them at 60°C for 27 hrs. We used a diamond knife to cut ultrathin sections. We observed the sections under a Philips CM12 transmission electron microscope.

### Quantification of DNA damage

We quantified the DNA damage caused by X-ray exposure by the silver-stained Comet alkaline assay (Travigen®, Cat#4251-050-K)^17,18^ according to the manufacturer’s specifications and we used ImageJ software^13^ to quantify the DNA fragmentation.

### Molecular genetic analysis

We treated the *T. wilhelma* specimens with 600 Gy (actually 518 Gy due to water absorbance). After 24 hours, 7 days, and 21 days following X-ray exposure we extracted the total RNA (RNeasy**^®^** mini kit, Qiagen, cat. n. 74104) from 3 sponges for each treatment and control. After verifying the purity and integrity of the RNA using an Agilent 2200 TapeStation, part of the extracted RNA (11.06 ng per sample on average) was utilized for RNA-seq analysis. We sequenced the samples using an Illumina NextSeq 500 instrument. We checked the quality of the RNA-seq reads for each sample using FastQC v0.10.1 and we aligned the reads with the reference genome (NCBI, SRA, SRR2163223) using STAR v2.5.1b (22.68 million reads uniquely mapped on average per sample). Cufflinks v2.2.1 was used to report FPKM values (Fragments Per Kilobase of transcripts per Million mapped reads) and read counts. We uniquely mapped 17.05 million reads to the reference genome on average, per sample. We performed a differential expression analysis using the EdgeR package from Bioconductor v3.2 in R 3.2.3. For each pairwise comparison, genes with false discovery rate (FDR) <0.05 were considered significant and 2 log_2_-fold changes of expression between conditions were reported after Bonferroni correction. We analyzed the differentially expressed genes using BLAST^19^, Ensembl^20^ and their functional annotations, including fold enrichment (FE), using DAVID^21,22^ and PANTHER^23^ software, and protein domains^24^. We focused on the overexpressed genes because the decrease of gene expression can be an nonspecific effect of X-ray exposure due to cellular damage.

## Results

We initially conducted a dose finding experiment to quantify the maximum tolerance to increasing doses (range:160-800 Gy) of radiation on sponges. After the treatment we observed the sponges daily. An 800 Gy dose is lethal for sponges (n=5, 80% lethality). In contrast, all sponges (n=7) exposed to 600 Gy suffered transitory morphological changes but survived. We then conducted the subsequent experiments using a single dose of 600 Gy.

### Morphological observations

We observed a general pattern in the morphological changes over time in sponges after X-ray exposure (Fig. 2, 3 and Fig. 1S).

a. Initially (2-7 days) the sponges begin to shrink, producing short and thin body extensions, and the pores and the osculum are no longer visible (Fig. 2, 3 and Fig. 1S). Sponges reach the minimum size after 21-25 days (Fig. 2, 3; paired t-test, df=16, t=6.341, p<0.0001), their surface became smooth and they produced large and long body projections, causing sponges to acquire an irregular star shape (Fig. 2, 3 and Fig. 1S).
b. Then, sponges reverse the shrinking process, but their morphologic features appear to be still altered or progress to further dissolvement (Fig. 2): body projections increased their surface and additional body projections were generated (Fig. 2). Body extensions were either re-absorbed or broke off, generating new satellite sponges, observed in 33.3% of sponges (Fig. 2-4 and Fig. 1S).
c. Sponges gradually reacquired their original anatomical organization and appeared normal after ∼180 days. Four out of 21 treated sponges died after an average of 162 (±30.9 S.D.) days. At the time of this writing, it has been over 1 year since treatment. 17 of 21 sponges treated with 600 Gy are alive, do not show any morphological changes, and are indistinguishable from untreated sponges (Fig. 2, 3 and Fig. 1S).

**Figure 2.**
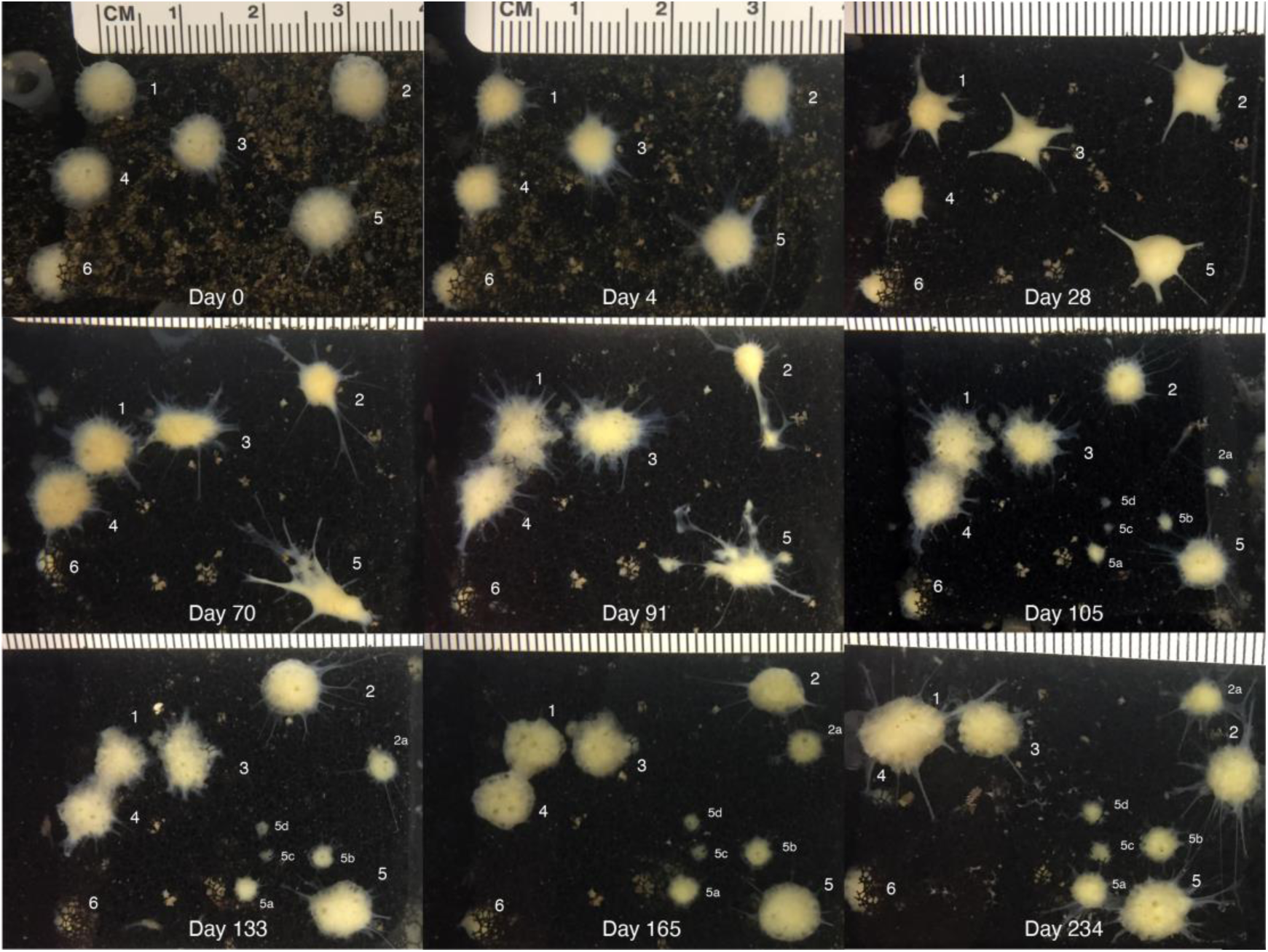
We observed morphological changes of 6 sponges after exposure to 600 Gy of X-rays. *T. wilhelma* sponges are capable of locomotion and their position can slowly change over time. The first picture was taken starting before the exposure (Day 0) and pictures were taken periodically until 234 days after the exposure. Each sponge is identified with a number. The offspring of the sponges are identified with the number of the parental sponge and a letter (e.g. sponge 2, offspring 2a). Sponge 5 had the most dramatic changes in morphology and generated 4 satellite sponges. Sponges 1 and 4 fused completely by day 234.

**Figure 3.**
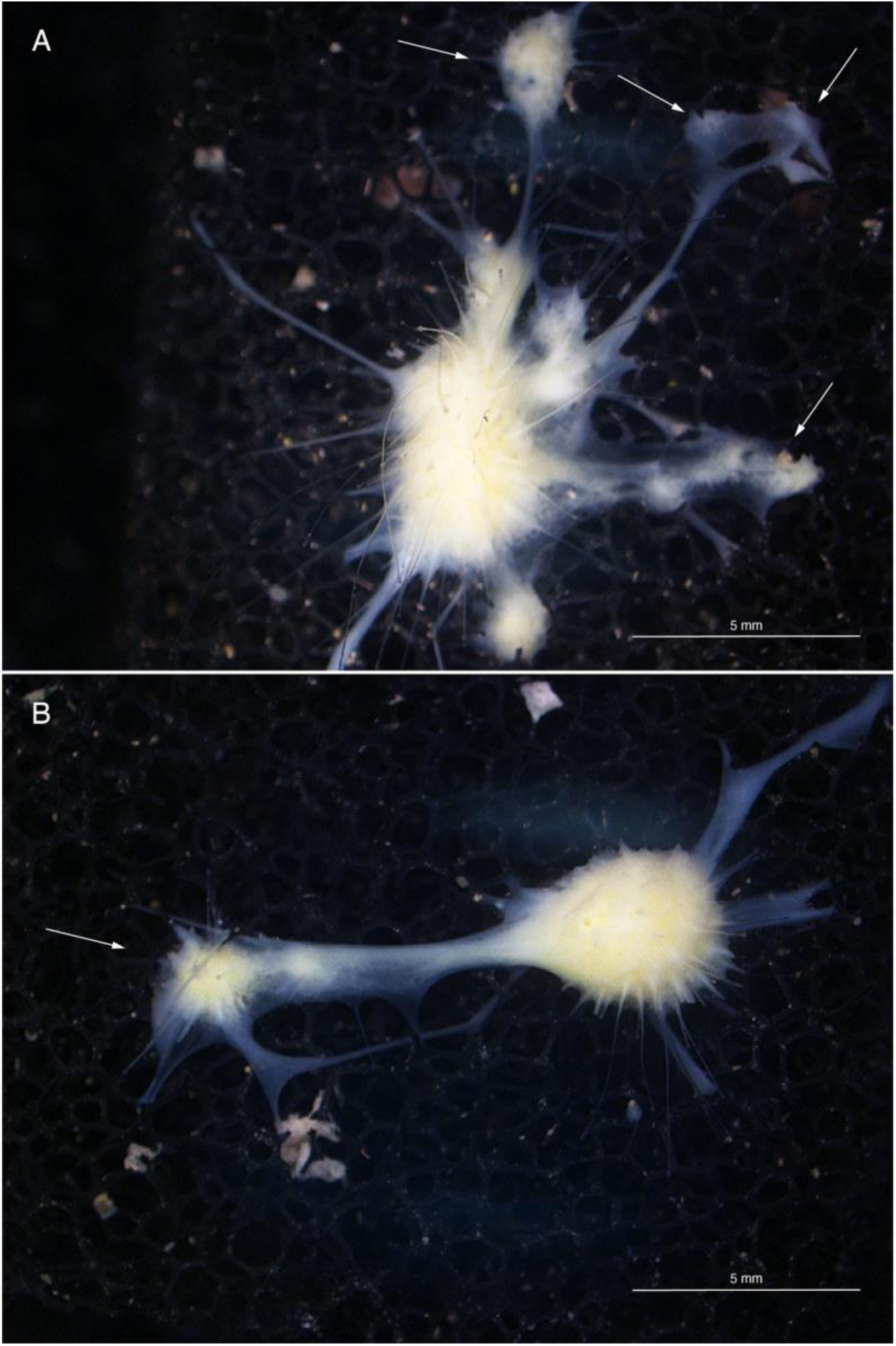
Morphologic changes in two sponges (A, B) 91 days after X-ray exposure. Sponges can develop extensive body projections. The body projections either generate new satellite sponges (arrows) or they are reabsorbed.

**Figure 4.**
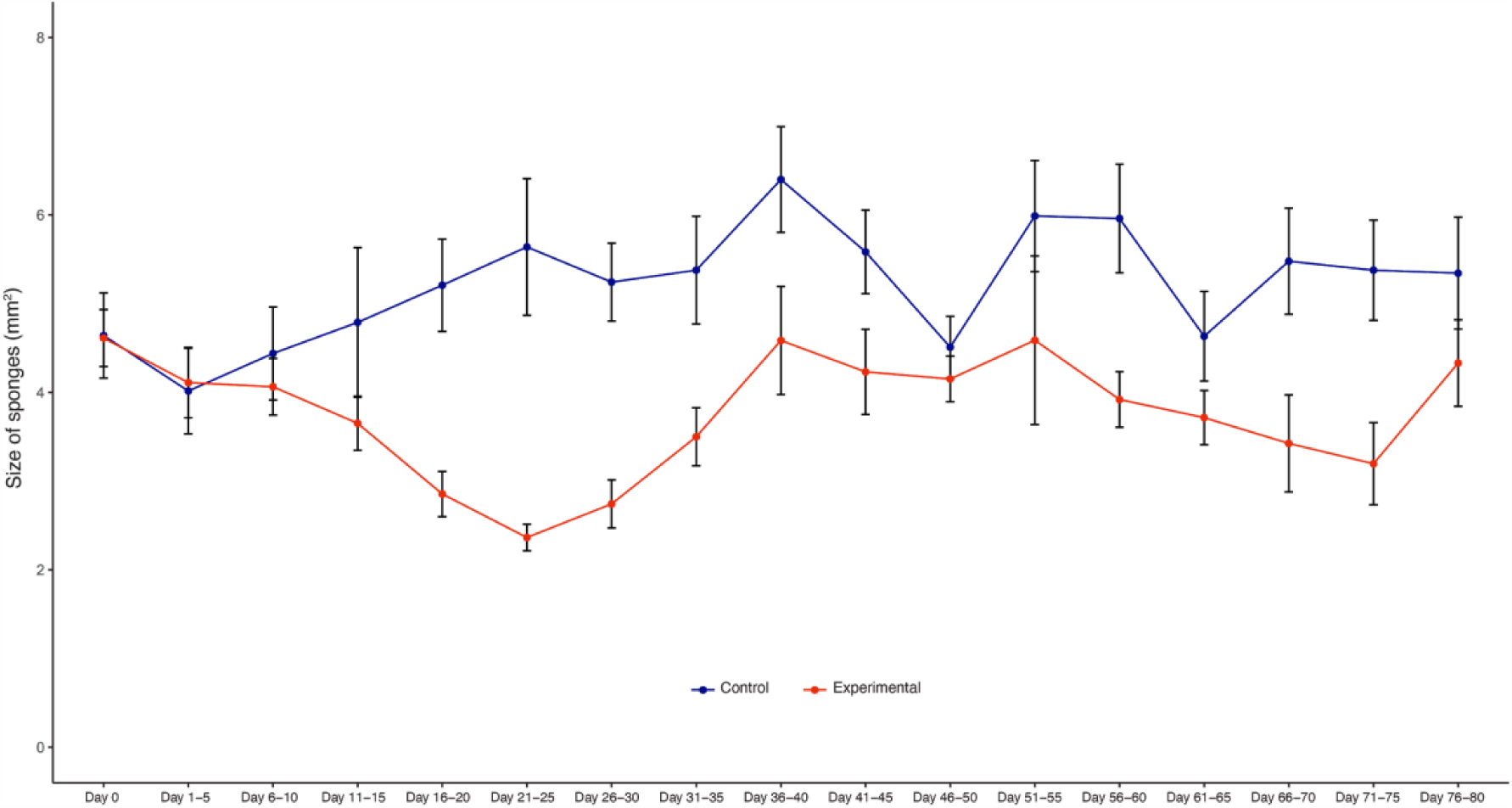
The size of sponges decreased after X-ray treatment (t-test controls vs X-ray treated sponges, paired by time point, df=16, t=6.341, p<0.0001). The blue line shows the control sponges (n=10) and the red line shows the X-ray treated sponges (n=21). Sponges reach the smallest size after 21-25 days. Then, they gradually increased in size and recovered their morphological features.

### Histological analysis

After X-ray exposure, sponges lose their typical anatomical organization^16^ (Fig. 5). The filtering structures of sponges (choanoderm) and the specialized water flowing and feeding cells (choanocytes) are lost (Fig. 5), and the choanoderm appear to be filled with undifferentiated cells^25^. The histological analyses did not show necrotic areas at any time after X-ray exposure (Fig. 5).

**Figure 5.**
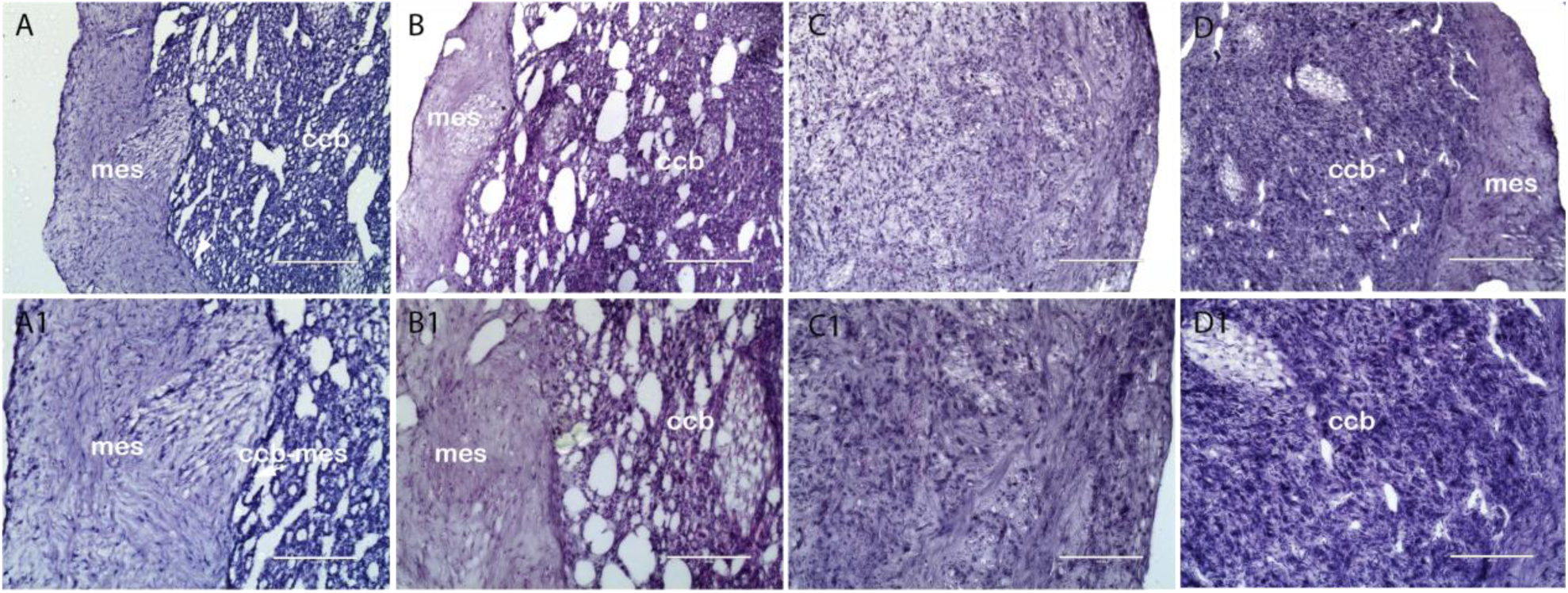
*T. wilhelma* untreated (A, A1), after 24 hours (B, B1), 7 days (C, C1), and 21 days (D, D1) from X-ray exposure. A1, B1, C1 and D1 (scale bar=100 μm) are a magnification of samples A, B, C, D and D (scale bar= 200 μm). Twenty-four hours after X-ray exposure, control sponges have anatomical features indistinguishable from untreated sponges (A vs. B). The normal anatomy is completely lost after 7 days and cells show an undifferentiated phenotype (C, C1). After 21 days most cells still appear to have an undifferentiated phenotype but anatomical reorganization has begun (Hematoxylin & eosin stain; ccb=choanoderm; mes=peripheral mesohyl; ccb-mes boundary= mesohyl of the cortex–choanoderm boundary).

### Electron microscopy analysis

We further investigated the morphological changes of the choanoderm after 7 days from X-ray exposure. Indeed, the choanoderm of treated sponges is disorganized and the choanocyte chambers are deeply altered or absent. The choanocytes are not anymore recognizable (Fig. 6). The mutualistic bacteria, presumably Cyanobacteria^26^, are able to survive the X-ray treatment as well.

**Figure 6.**
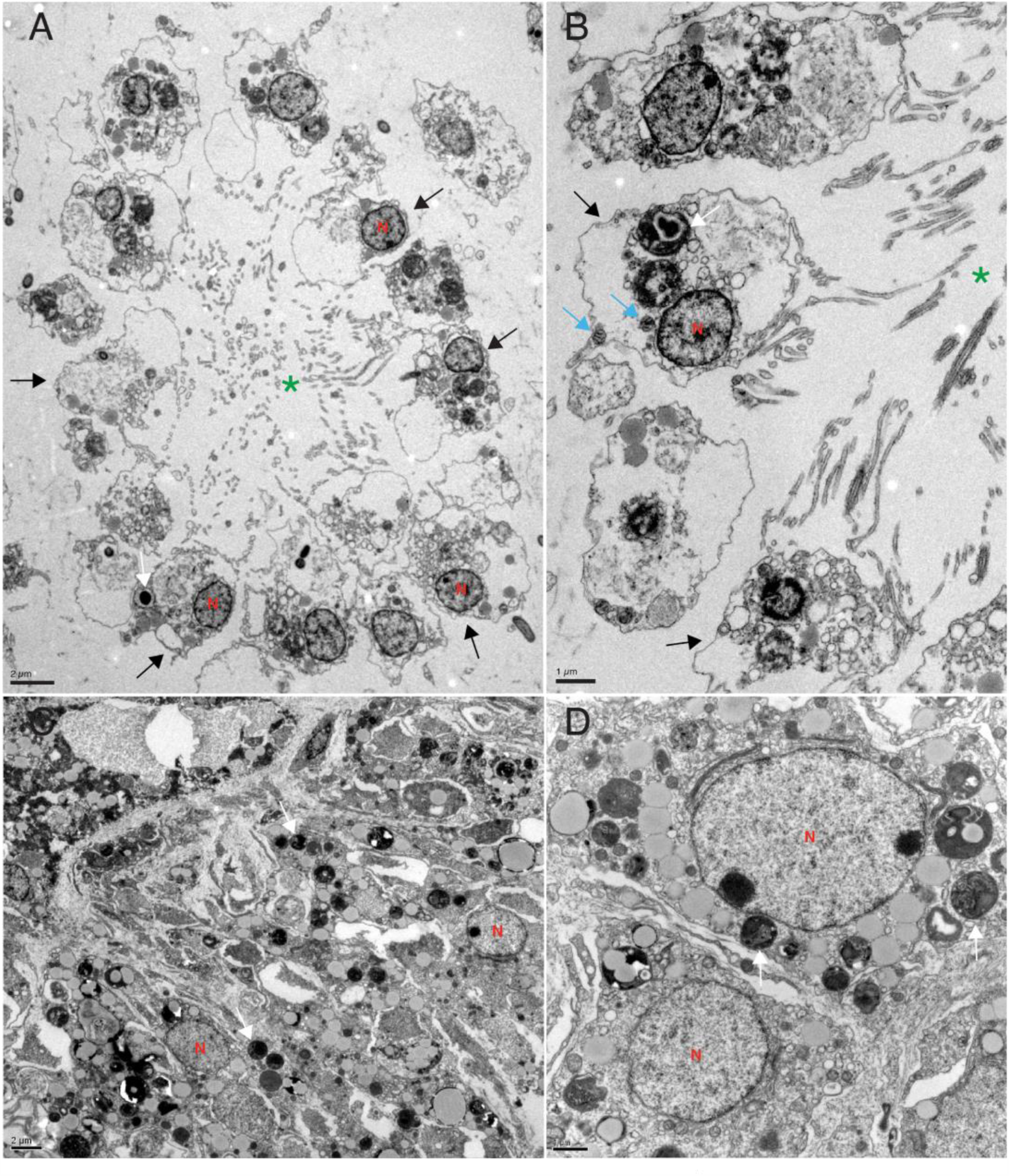
Transmission electron microscopy analysis of control (A, B) and X-ray exposed sponges (C-F), 7 days after exposure. The choanoderm of control sponges is well organized with choanocyte chambers (green asterisk *). Instead, the choanoderm of treated sponges (C-F) is disorganized and the choanocyte chambers are absent. The X-ray exposed choanoderm is characterized by the extensive presence of vesicle and membrane heterogeneous structures. The nuclei appear to be intact and have different shapes. Black arrows=choanocytes, blue head arrow=mitochondria, N=nucleus, white arrows=bacteria.

### DNA damage analysis

DNA fragmentation analysis (Comet assay) showed limited DNA degradation immediately after a submaximal (600 Gy) X-ray exposure (DNA fragmentation, treated: 8.23% ± 16.32 S.D., controls 1.34% ± 6.99 S.D. (Fig. 7).

**Figure 7.**
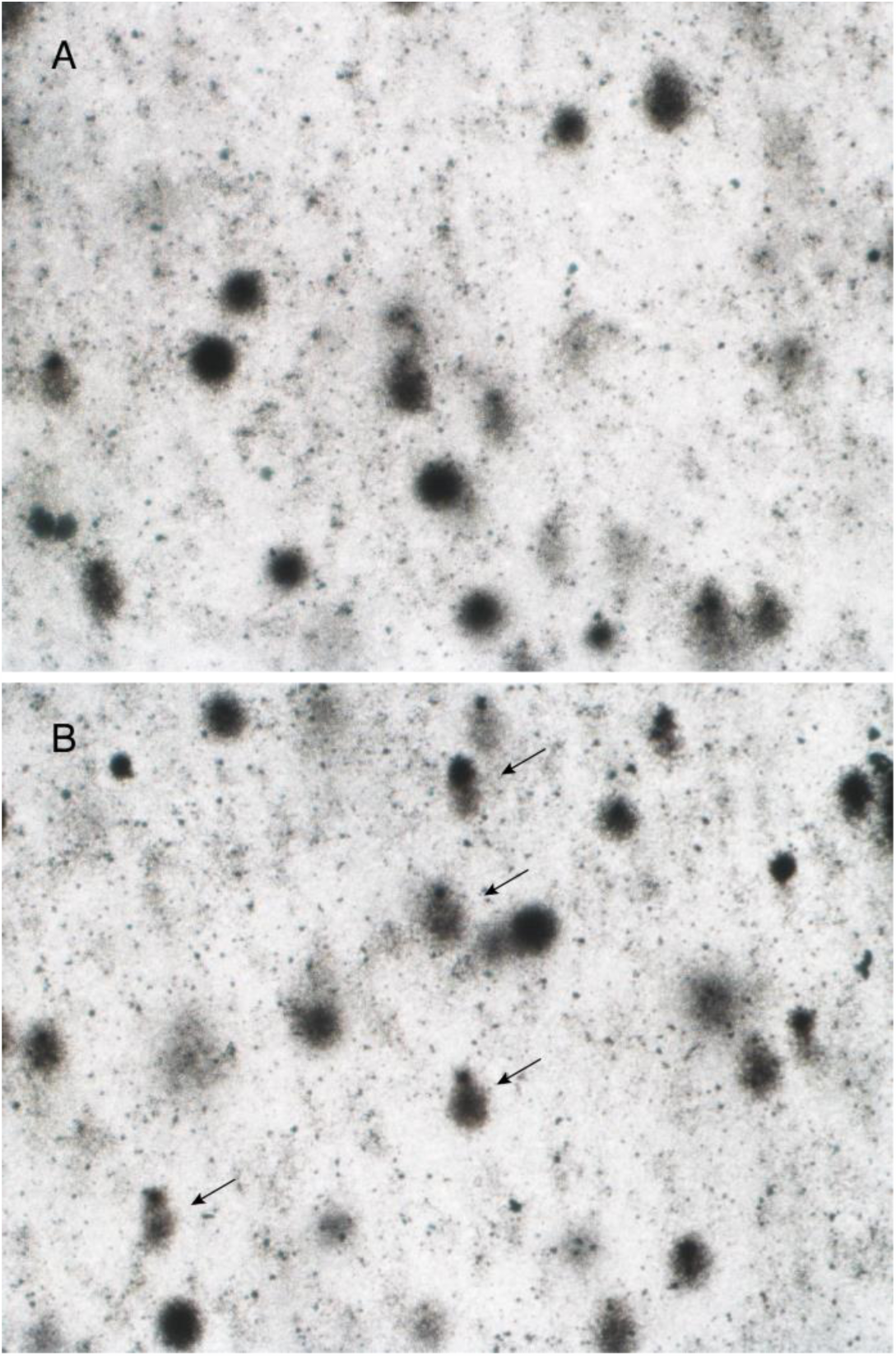
The X-ray treatment produces little DNA fragmentation as measured by the Comet assay. There is evidence of DNA in: 8.23% ± 16.32 S.D. of treated sponges, and 1.34% ± 6.99 S.D. controls. Representative images of control (A) and X-ray treated (B) sponge nuclei, arrows indicate examples of partially fragmented nuclei (comets).

### Gene expression analysis

We performed the transcriptome analysis (RNA-seq) of 3 *T. wilhelma* specimens collected at each of 3 different time points (24 hours, 7 and 21 days) after X-ray treatment. We found a total of 639 overexpressed transcripts in the three experimental time points compared to untreated sponges at the same time points (Fig. 8). Each group had different gene expression levels and differs from the controls (Fig. 2S). Many of the expressed genes have a human homolog (given in parentheses). There are genes overexpressed only at a specific time point (24 hours, 7 or 21 days) and genes overexpressed at 2 or 3 time points (Tab. 3S, fig. 8 and fig. 3S).

**Figure 8.**
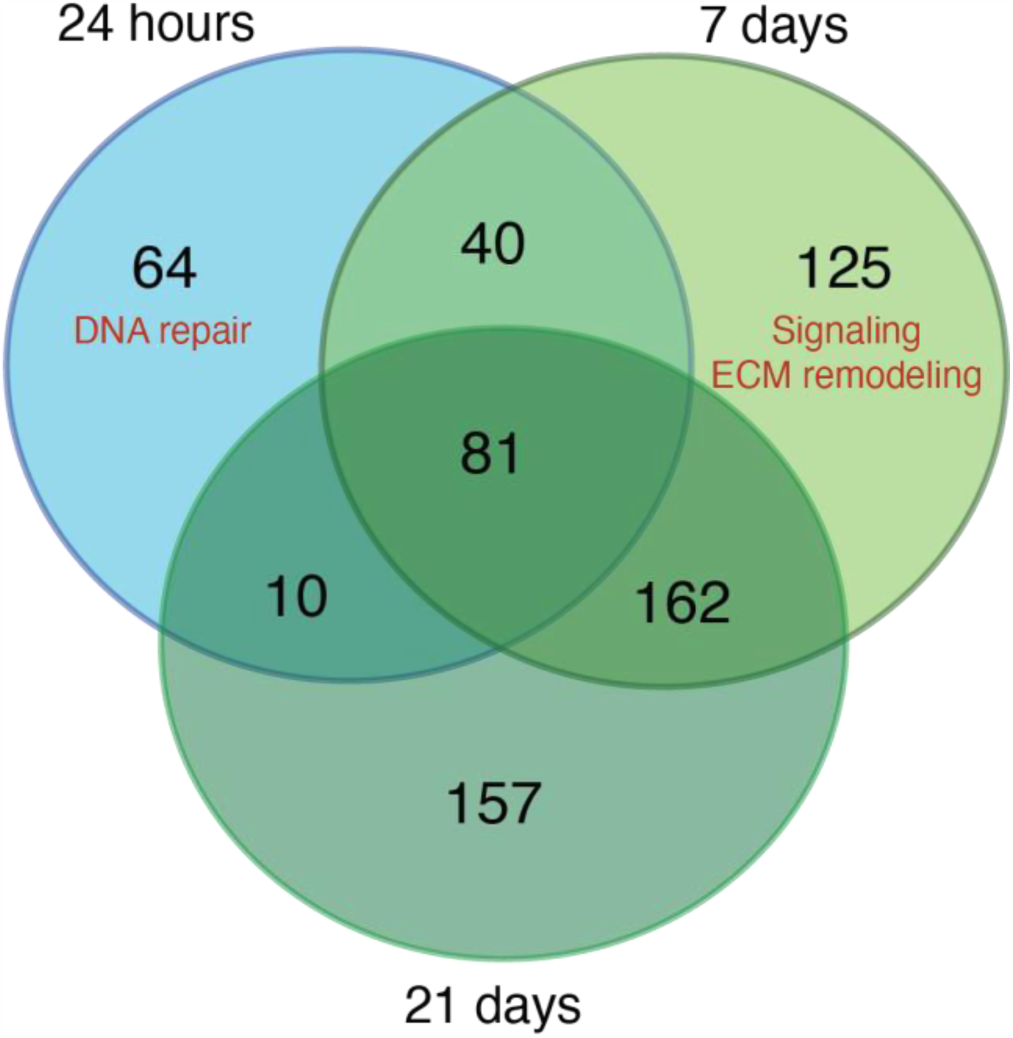
Venn diagram shows the number of genes that were over-expressed at the 3 different time points (24 hours, 7 and 21 days) after X-ray treatment. There are genes overexpressed only at a specific time point and genes overexpressed at 2 or 3 time points. The number of transcripts detected increased from 24 hours to 21 days. 24 hours and 21 days share only 1.6% of transcripts, on the contrary 7 days and 21 days have the largest percentage (25.4%) of overlapping transcripts. Genes can be activated at different times and for a different length of time after X-ray exposure. Venn diagram shows the exact number of transcripts expressed. There is an enrichment of genes involved in DNA repair specifically expressed after 24 hours and in signaling after 7 days. EMC= extracellular matrix.

#### Genes overexpressed 24 hours after X-ray exposure

We found 195 transcripts overexpressed 24 hours after X-ray exposure, of which 97 have a human homolog (Fig. 8, tab. 1S and fig. 3S). Sixty-four of those 195 transcripts were only overexpressed after 24 hours but not at later time points. Fourty-four of those 64 overexpressed transcripts specific to the 24-hour time point have a human homolog. We detected an enrichment of human homolog genes involved in DNA repair (FE=10.3, FDR= 0.037, DAVID) such as twi_ss.22376.1 (*LIG3*), DNA ligase and twi_ss.21448.1 (*MRE11*), involved in the double strand break repair mechanisms, twi_ss.11375.1 (*PCNA*), proliferating cell nuclear antigen, has a key role in DNA damage response, twi_ss.28211.1 (*RPA1*), DNA repair pathway and the replication protein A1, is a cofactor of DNA polymerase delta involved in the RAD6-dependent DNA repair pathway, Twi_ss.25822.1-3 (*GINS4*), GINS subunit, domain A, is involved in double-strand break repair via break-induced replication.

We found that twi_ss.17326.7 gene (*CUBN*), cubilin, is the most differentially expressed gene (logFC=8.55) specific to the 24-hour time point after X-ray exposure. *CUBN* is an endocytic receptor expressed in the epithelium of intestines and kidneys^27^, and down-regulated in renal cell carcinoma^28^.

In addition, we identified overexpression of genes involved in stress response such as twi_ss.28284.1 (*HSPA1A*), the heat shock protein family A (Hsp70) member 1A.

#### Genes overexpressed 7 days after X-ray exposure

We found 408 transcripts overexpressed, of which 141 have a human homolog (Fig. 8, tab. 1S and fig. 3S). Among them, 125 transcripts were specifically overexpressed after 7 days, of which 53 have a human homolog. We detected an enrichment of ankyrin containing domain genes (FE=19, FDR< 0.001), signal proteins (FE=2.4, FDR= 0.005) and extracellular matrix remodeling genes (FE=68.1, FDR= 0.03).

We identified genes specifically overexpressed after 7 days involved in DNA repair such as twi_ss.10792.1 (*FAN1*), twi_ss.27976.3 (*KAT5*) and twi_ss.19635.1 (*TRIP12*) and, importantly, we found a correspondence with the morphological changes observed 7 days after X-ray exposure and the function of genes overexpressed at the same time. For instance, we found the overexpression of genes involved in development, adult tissue homeostasis and mesenchymal transition such as twi_ss.2267.1 (*NOTCH1*), twi_ss.5628.6 (*MET*) MET proto-oncogene, receptor tyrosine kinase, twi_ss.6204.1 (*POSTN*), periostin. We also found genes involved in embryonic stem cell regulation such as twi_ss.7082.1 (*ETV4*), twi_ss.4355.1 (*TRIM71*), twi_ss.329.1 (*ZFP36L1*) together with a variety of genes such as twi_ss.26378.1 (*COL6A3*) collagen type VI alpha 3 chain, twi_ss.31853a.1 (*COL6A6*) collagen type VI alpha 6 chain, twi_ss.25970.2 (*LTBP1*) latent transforming growth factor beta binding protein 1, twi_ss.20516a.5 (*SORL1*) sortilin related receptor 1, twi_ss.2619.7 (*VWF*) von Willebrand factor, twi_ss.19885.1 (*NRXN3*) neurexin 3, twi_ss.1244.5 (*ADGRE5*) adhesion G protein-coupled receptor E5 involved with the epithelial-mesenchymal transition.

#### Gene overexpressed 21 days after X-ray exposure

we found genes 410 overexpressed transcripts of which 150 of these have a human homolog (Fig. 8, tab. 1S and fig. 3S). We identified 157 transcripts specifically overexpressed after 21 days of which 72 have a human homolog. There is not a functional signature specific to the genes expressed specifically after 21 days. We found genes involved in DNA double-strand break repair: twi_ss.10068.1 (*PARP3*) poly (ADP-ribose) polymerase family member 3, twi_ss.28718.1 (*PARPF19*) PHD finger protein 19 and, DNA repair: twi_ss.7757.3 (*UBR5*) ubiquitin protein ligase E3 component n-recognin 5.

#### Gene overexpressed 24 hours, 7 and 21 days after X-ray exposure

We found 81 transcripts overexpressed at all 3 time points of which 34 have a human homolog (Fig. 8, tab. 1S and fig. 3S). For instance, Twi_ss.16656.2 (*PHF8*), PHD Finger Protein 8 one of the most differentially expressed gene (9.3±1.8 S.D. logFC), the *C. elegans* homolog promotes DNA repair via homologous recombination^29^. Twi_ss.4977.9 (8.9±0.4 S.D. logFC) is a homolog of the human gene Spatacsin (*SPG11*). The function of *SPG11* is not well understood. It has a role in a form of spastic paraplegia, a neurodegenerative disorder and it appears to be also involved in DNA repair^30^.

#### Gene overexpressed 24 hours and 7 days after X-ray exposure

we found 40 transcripts overexpressed at both time points of which 19 of these genes have a human homolog (Fig. 7-9, tab. 1S). The most differential expressed gene (8±1.8 S.D. logFC) is twi_ss.12458.1 (*TTN*), a gene unknown to be activated after X-ray exposure.

#### Gene overexpressed 24 hours and 21 days after X-ray exposure

we found only 10 overexpressed transcripts of which 5 of these genes have a human homolog gene (Fig. 8, tab. 1S and fig. 3S). With only 5 genes with known homolog functions, there were not statistically significantly enriched pathways. However, twi_ss.19378.2 (*ERCC1*) is involved in DNA repair.

#### Gene overexpressed 7 days and 21 days after X-ray exposure

we found genes 162 overexpressed transcripts of which 45 of these have a human homolog gene (Fig. 8, tab. 1S and fig. 3S). Overall, there is an enrichment of genes involved in extracellular matrix organization such as fibronectin (FE=15.7, FDR< 0.0001), with an EGF-like domain (FE=19.7, FDR<0.0001) and signal peptides (FE=3.3, FDR<0.0001). One of the most overexpressed genes (10.1±0.7 S.D. logFC) is twi_ss.21105.9 (*SCUBE1*) signal peptide, CUB and EGF-like domain-containing protein 1, which may function as an adhesive molecule^31^.

## Discussion

*T. wilhelma* sponges can withstand 600 Gy (actual 517.6 Gy) of X-ray radiation. That is approximately 60 times the lethal dose for mice^32,33^ and 100 times the lethal dose for humans^34^. According to conventional wisdom, this amount of radiation should shatter the sponge’s DNA, however the Comet assay suggests this does not happen in *T. wilhelma*.

Early organisms evolved in an environment with higher levels of background radiation^35^. Despite the fact that water partially shields aquatic organisms from direct radiation exposure, radionuclides can accumulate in the sea. As filter-feeding animals, sponges could be particularly exposed to the accumulation of radionuclides and other toxic agents^36^. Moreover, sponges are sessile organisms without a nervous system thus not capable of rapidly escaping or quickly reducing the water flow through their bodies if the concentration of radioactive agents increases. For these reasons sponges are considered biological indicators of environmental pollution such as radionucleotides^37,38^.

There are examples of extreme radioresistance in bacteria^39^ and multicellular organisms capable of anhydrobiosis (desiccation) such as rotifers^40^ and tardigrades^41–43^, which is unlikely to apply to *T. wilhelma*. Tolerance for both desiccation and radiation may originate from similar DNA protective or repair mechanisms^40,44^. Indeed, desiccation causes DNA breakage^40,42^, similar to damage induced by radiation, that may be repaired upon rehydration^40^. Tardigrades possess molecular mechanisms to prevent DNA damage. Dsup is a tardigrade-specific nucleosome-binding protein that protects chromatin from hydroxyl radicals and contributes to the organism’s radio-tolerance^45^. The combination of DNA protective and DNA repair mechanisms determines the level of radioresistance of these organisms. Importantly, rotifers and tardigrades have no or highly restricted somatic cell turnover^41,46,47^, and the species tested for radioresistance have a short (∼60 days) lifespan^41,48^ which prevents mutant clones and accumulating further mutations, leading to cancer. In these conditions, the DNA damage that occurs does not propagate and so might not be apparent^49^. Thus, there is little chance for cancer to develop in tardigrades.

In order to investigate the long-term effect of DNA damage and cancer in invertebrates, somatic cell turnover and long lifespan should be an essential feature of any experimental organism. However, the main invertebrate model organisms currently in use (e.g. *Caenorhabditis elegans* and *Drosophila melanogaster*) have limited or no somatic cell turnover^50^ and their lifespans^51,52^ are too short to study the long term effects of radiation. In contrast, sponges have somatic cell turnover^2^ and a remarkably long lifespan: the largest *Xestospongia muta* specimen described on Caribbean reefs is estimated to be more than 2,300 years old^53^, a specimen of the sponge *Monorhaphis chuni* is thought to be 11,000 years old^54^ and radiocarbon dating of the sponge *Rossella racovitzae racovitzae* determined that was around 440 years old^55^. Selection for this long lifespan may have also selected for cancer suppression mechanisms.

Considering all these factors: somatic cell turnover, long life span and a primitive immune system^56^, we would expect the development of tumors in sponges, but there have been no reports of cancer in the entire Porifera phylum^1^ (with 8500 described living species^57^).

In our experimental setting, (single high dose) radiation did not induce cancer development in *T. wilhelma* during 1 year following X-ray exposure. Malignant cancer risk in humans is estimated to be 8% per Gy^58^. A small fraction of the dose to which the sponges have been subjected would generally have given rise to cancer in mice^59^ and humans^58^.

The molecular data and morphological observations suggest that sponges protect their DNA from damage in the first place and then activate mechanisms of DNA repair and cell death as they go through a complex phase of tissue reorganization. Finally, they rebuild their tissues’ original features. Our findings suggest that the cancer resistance in sponges might be linked to their radioresistance.

Genes involved in DNA repair are activated at different times and for different lengths of time, providing insight into the temporal activation of these genes during the DNA repair process (Fig. 8, tab. 1S and fig. 3S). We found the overexpression of genes known to have a role in DNA double strand break repair (Tab. 1S), confirming that our experimental setting is capable of induced the typical DNA damage produced by X-ray exposure and that sponges respond to radiation by upregulating DNA repair. As expected, we observed a higher number of genes involved DNA repair 24 hours and 7 days after X-ray exposure, but we also identified genes involved in this process specifically expressed 21 days after X-ray exposure. This observation suggests that after 21 days the sponges are still actively repairing the DNA damage induced by radiation. Treated sponges overexpressed only 81 transcripts (12.7% of all overexpressed transcripts) in all 3 time points, but overexpressed 64 transcripts only at 24 hours, 125 transcripts only at 7 days, and 157 transcripts only at 21 days, suggesting waves of sequential transcriptional events.

We found overexpressed genes previously not linked to X-ray induced damage, or with unknown function in humans (Table 3S). For example, twi_ss.17326.7 (*CUBN*), cubilin in humans. CUBN is an endocytic receptor expressed in the epithelium of intestines and kidneys^27^. Interestingly, low expression of *CUBN* in renal cell carcinoma is significantly associated with early disease progression and poor patient outcome^28^. We hypothesize that the *CUBN* gene has a protective function against DNA damage or is involved in DNA repair and thus its downregulation in kidney cancer increases the chance of an aggressive evolution of the disease.

Epigenetic changes such as methylation are induced by X-ray exposure and contribute to regulate the cell response to stress^60^. Though we did not directly measure methylation, we detected the overexpression of twi_ss.21704.3 (*NSUN7*), a gene involved in methylation. Methylation induced by radiation can be a persistent epigenetic change after radiation ^60^ regulating gene activity long after expression normalization.

In addition to mechanisms of DNA repair, we detected the overexpression of genes involved in apoptosis and cell death. For example, Twi_ss.31154.2 (*NOX5*), NADPH oxidase–generated ROS, a gene involved in heavy ion irradiation–induced cell death^60^, suggesting the activation mechanisms of cell death, presumably to remove cells too compromised by radiation.

Notably, we observed a distinct pattern in the morphological changes in the treated sponges. Seven days after X-ray exposure, sponges lose their typical anatomical organization and most of the cells seem to acquire undifferentiated features. Totipotent cells (archeocytes) are part of the mesohyl and they can replace damaged cells^61^. Choanocytes, one of the most specialized types of sponge cells, can also transdifferentiate into archaeocytes (stem cells) and serve the same function^61,62^. In transcriptomic analysis 7 days following X-ray exposure, we found overexpression of genes known to be involved in cell adhesion, signaling, embryonic cell regulation and EMT such as *NOTCH1* and *MET*, consistent with our observations of changes in sponge morphology.

### Studying radiation resistance in sponges might help improve radiation therapy

Ionizing radiation is part of the natural environment and evolution of organisms. Thus, the study of radiation responses in sponges could contribute to understanding the molecular evolution of cell radioresistance. This could help us understand the evolution of radioresistance in cancer cells as well, and potentially lead to methods for protecting human cells from radiation damage.

Over the course of a human lifetime, cells can evolve mechanisms that allow them to endure the effects of X-ray exposure^63,64^ through somatic evolution. In human cancers, radiotherapy often induces EMT, and leads to increased radioresistance accompanied by increased cell migration and invasion^65–68^.

The finding that sponges react to X-ray exposure by increasing the number of undifferentiated cells suggests that EMT is not a cancer cell specific response to radiotherapy, but rather a natural cellular response to radiation both in organisms and human cells^69^. Sponges are particularly suitable to study this process because their simple tissue organization facilitates identification and analysis of tissue alterations. Sponges give us the unique opportunity to study mesenchymal proliferation and EMT in response to radiation, removing the confounding effect of thousands of mutations or deregulated genes described in cancers which may be unrelated to EMT. It is important to note that cancer cells do not utilize *de novo* cancer specific mechanisms to survive radiation exposure, but rather employ a biological process that is shared with sponges and has been evolutionarily conserved in our genomes for 100’s of millions of years. Understanding this biological process in response to radiotherapy in sponges could provide the basis for new treatment and cancer prevention strategies.

The study of the evolution of anti-cancer mechanisms such as radiation resistance can contribute to the understanding of multicellular organisms more generally, as the evolution of multicellularity itself depended on the multicellular organisms’ ability to prevent somatic cells proliferating out of control^70^. Our work suggests that sponges may be particularly resistant to cancer because of their radiation resistance, and shows that sponges are an viable model system for studying anti-cancer mechanisms and radiation resistance.

## Acknowledgments

We would like to thank Erik Southard for helping to maintain the sponges and assistance during the experiments; Joy Blain and Shanshan Yang (ASU Genomics Facility) for RNA sequencing. David Lowry (Life Science Electron Microscopy Lab) for TEM imaging; Debra Baluch (W.M. Keck Bioimaging Laboratory) for histological samples preparation for light microscopy. We are also grateful to Tim Chan for his advice and suggestions. This work was supported in part by NIH grants U54 CA217376, U2C CA233254, P01 CA91955, R01 CA170595, R01 CA185138 and R01 CA140657 as well as CDMRP Breast Cancer Research Program Award BC132057 and the Arizona Biomedical Research Commission grant ADHS18-198847. The findings, opinions and recommendations expressed here are those of the authors and not necessarily those of the universities where the research was performed or the National Institutes of Health.

## Author contributions

A.F, A.A. and C.C.M. designed the study. A.F. designed and performed the experiments, collected and analyzed the data. J.T. was an undergraduate student, he contributed to perform the experiments and collecting data. J.S. is an undergraduate student, he contributed to perform the bioinformatic analysis. A. Fortunato, A.A. and C.C.M. wrote the manuscript.

---

**Figure supplementary 1S.**
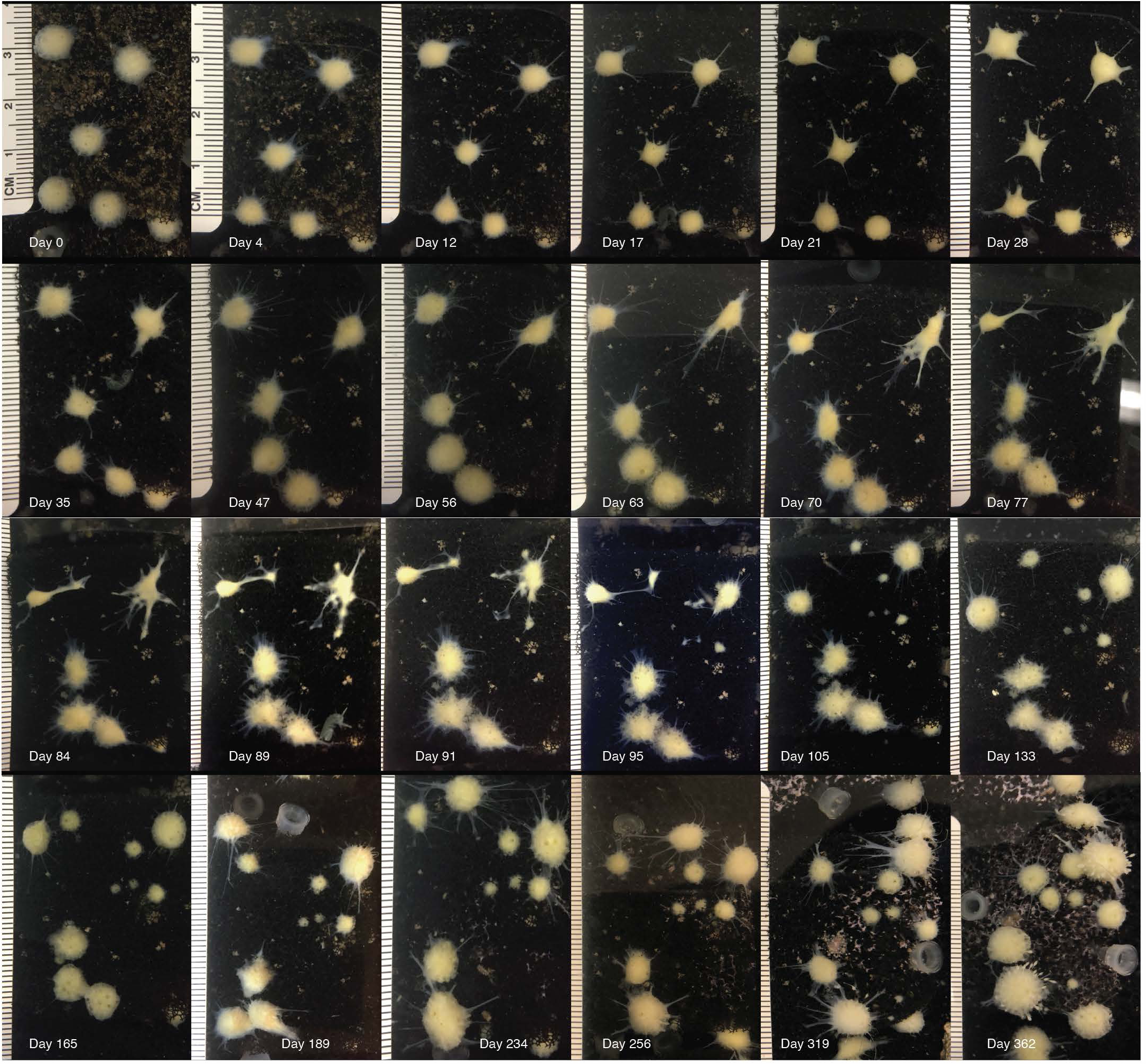
Long term observation of sponge’s morphology. Initially sponges suffer acute morphological changes but they recover from the X-ray treatment. Images of the same group of sponges from o to 362 days after X-ray exposure.

**Figure supplementary 2S.**
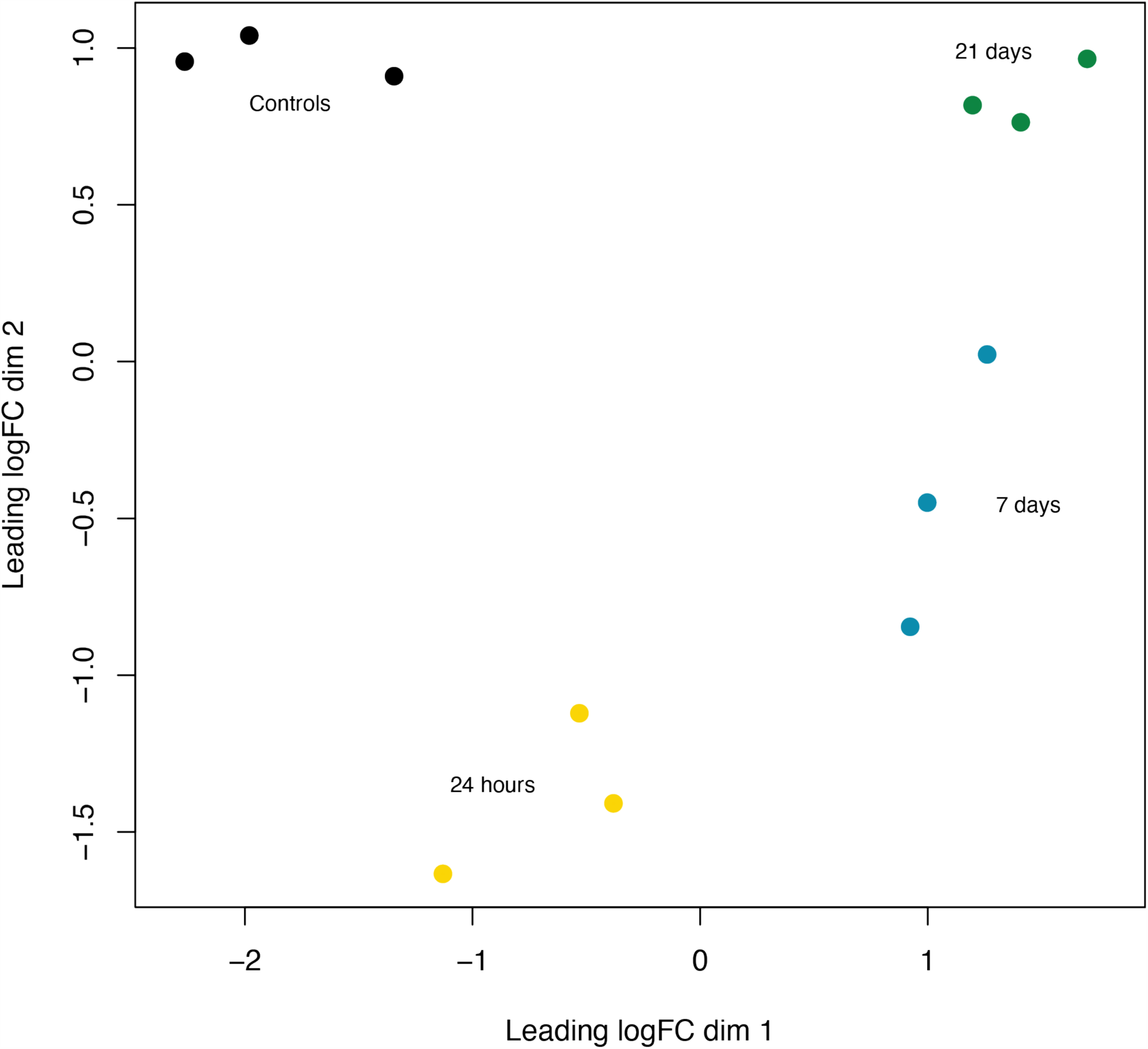
Multidimensional scaling (MDS) plot. There is a gene expression similarity between the samples collected at the same time point and a gene expression distance between the groups: controls, 24 hours, 7 and 21 days.

**Figure supplementary 3S.**
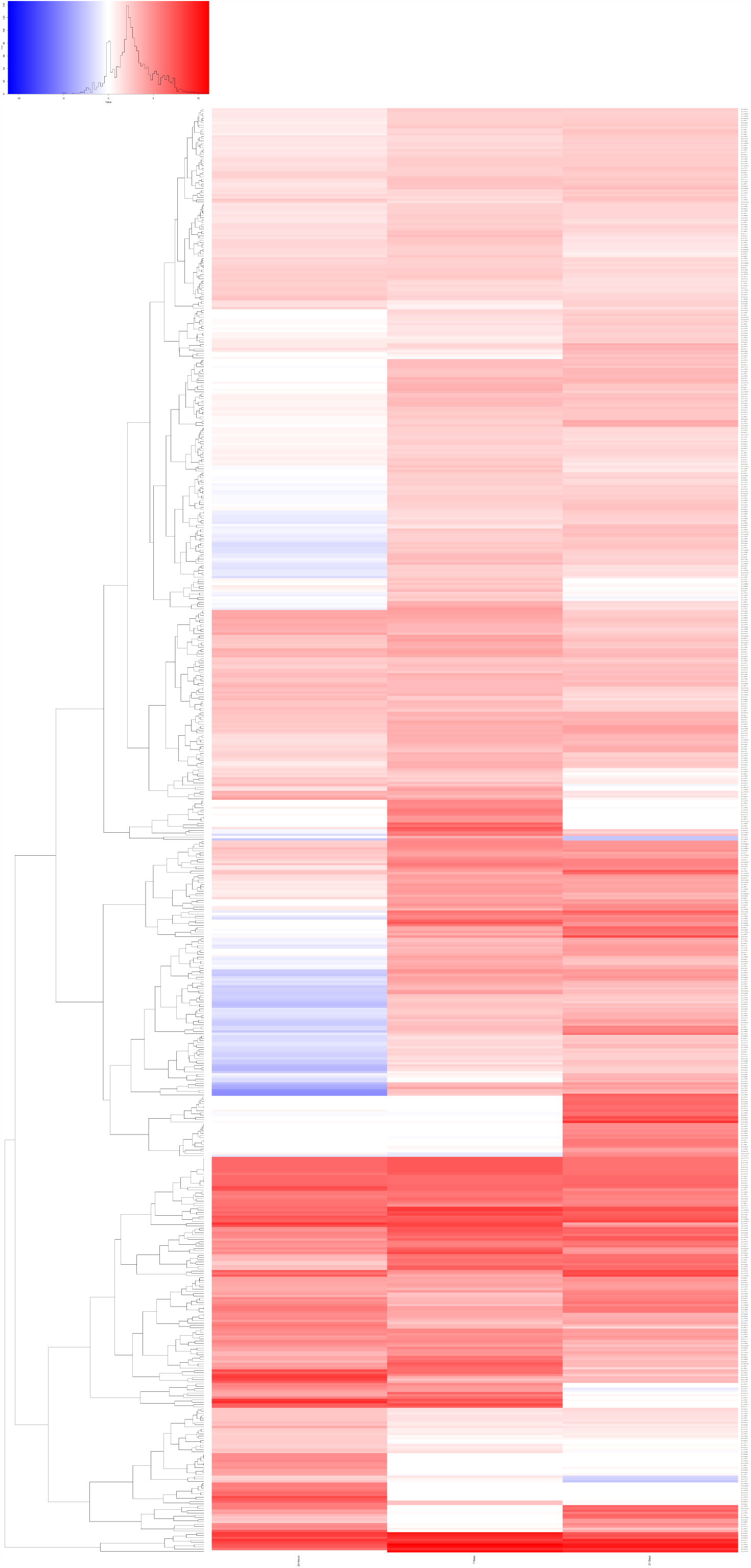
Heatmap of RNA-seq analysis. There are sequential transcriptional events following X-ray exposure. Genes can be expressed only to a specific time point, 24 hours, 7 or 21 days or at 2 or 3 time points.

**Table 1S.**
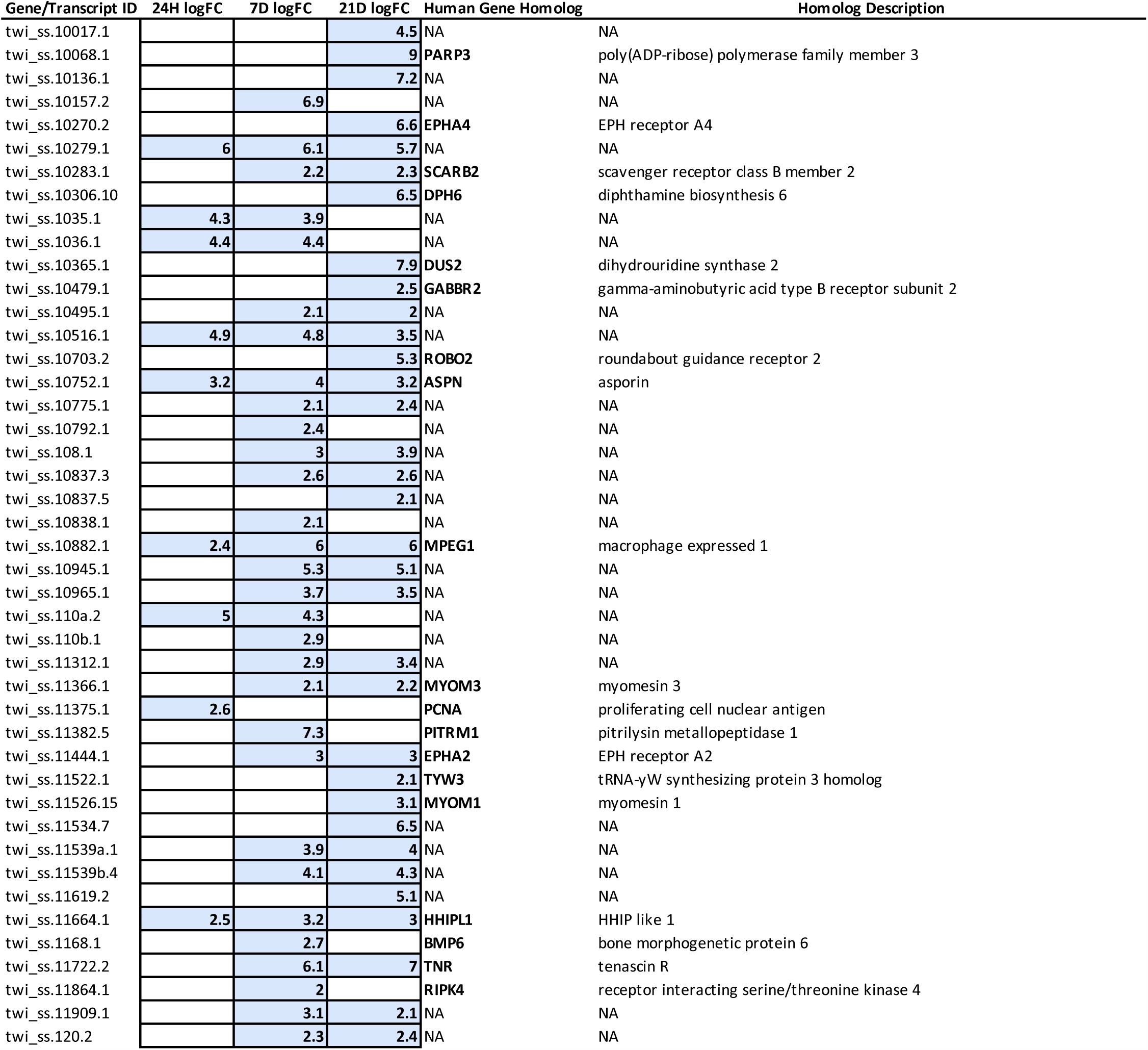
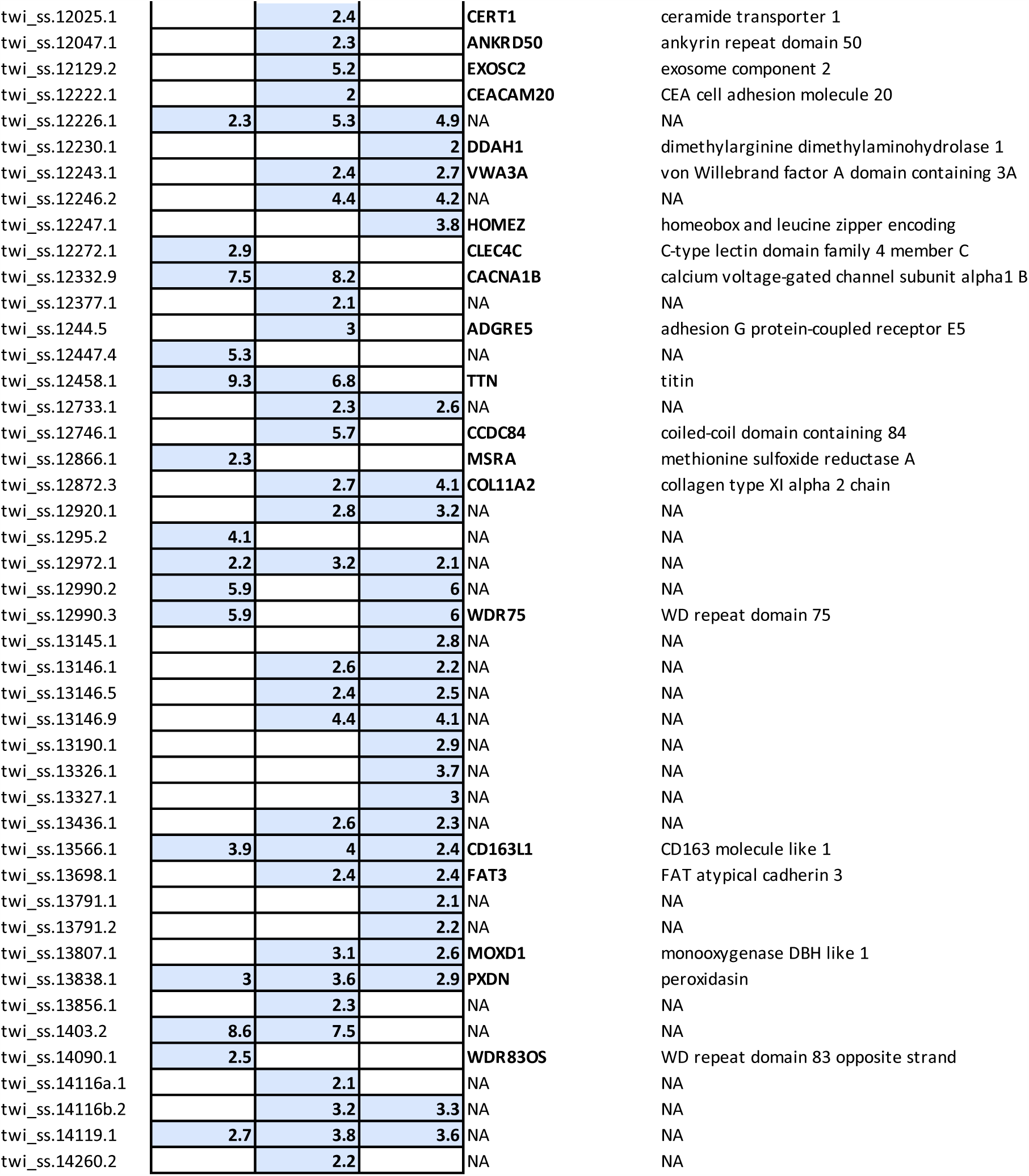
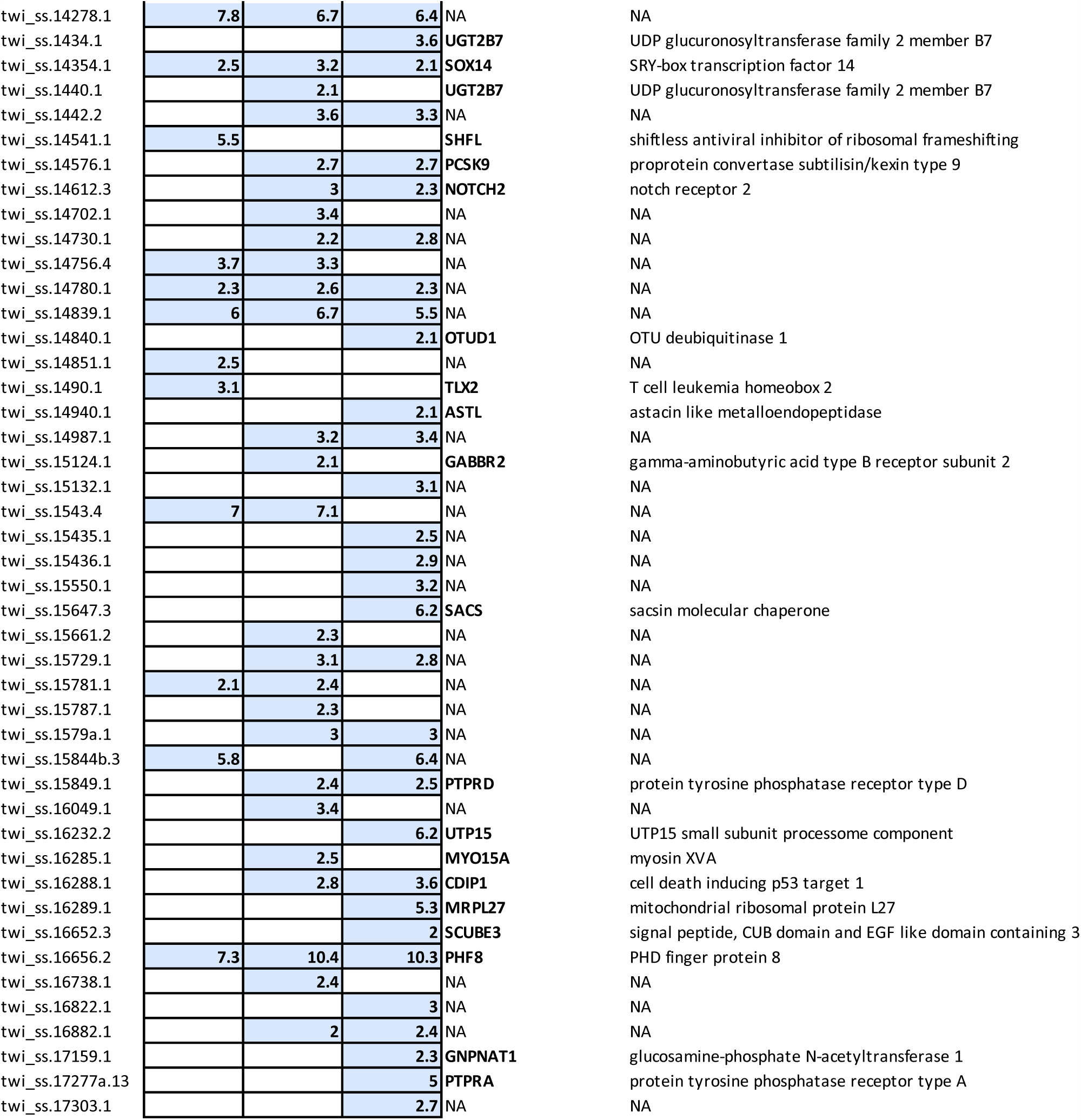
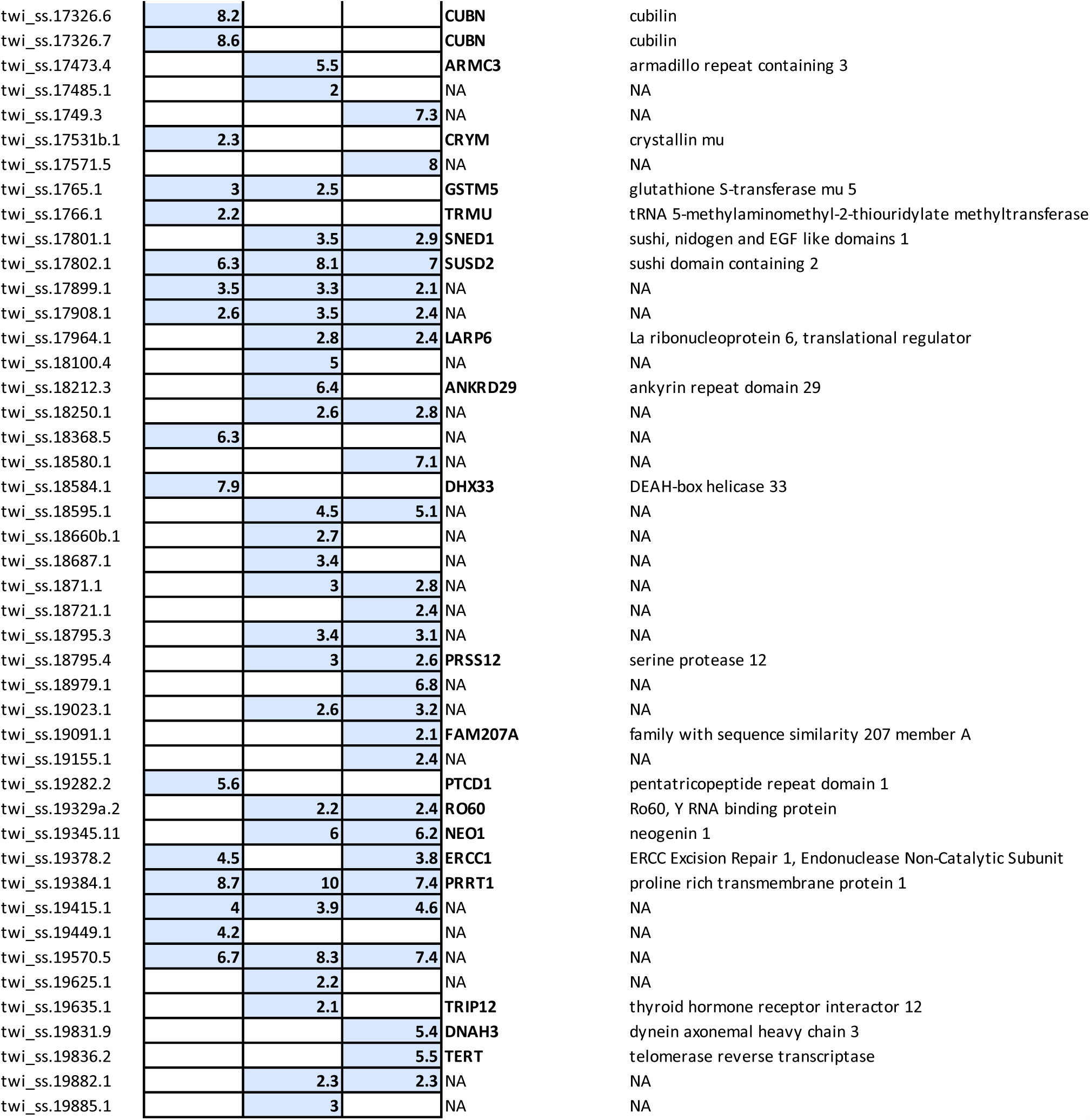
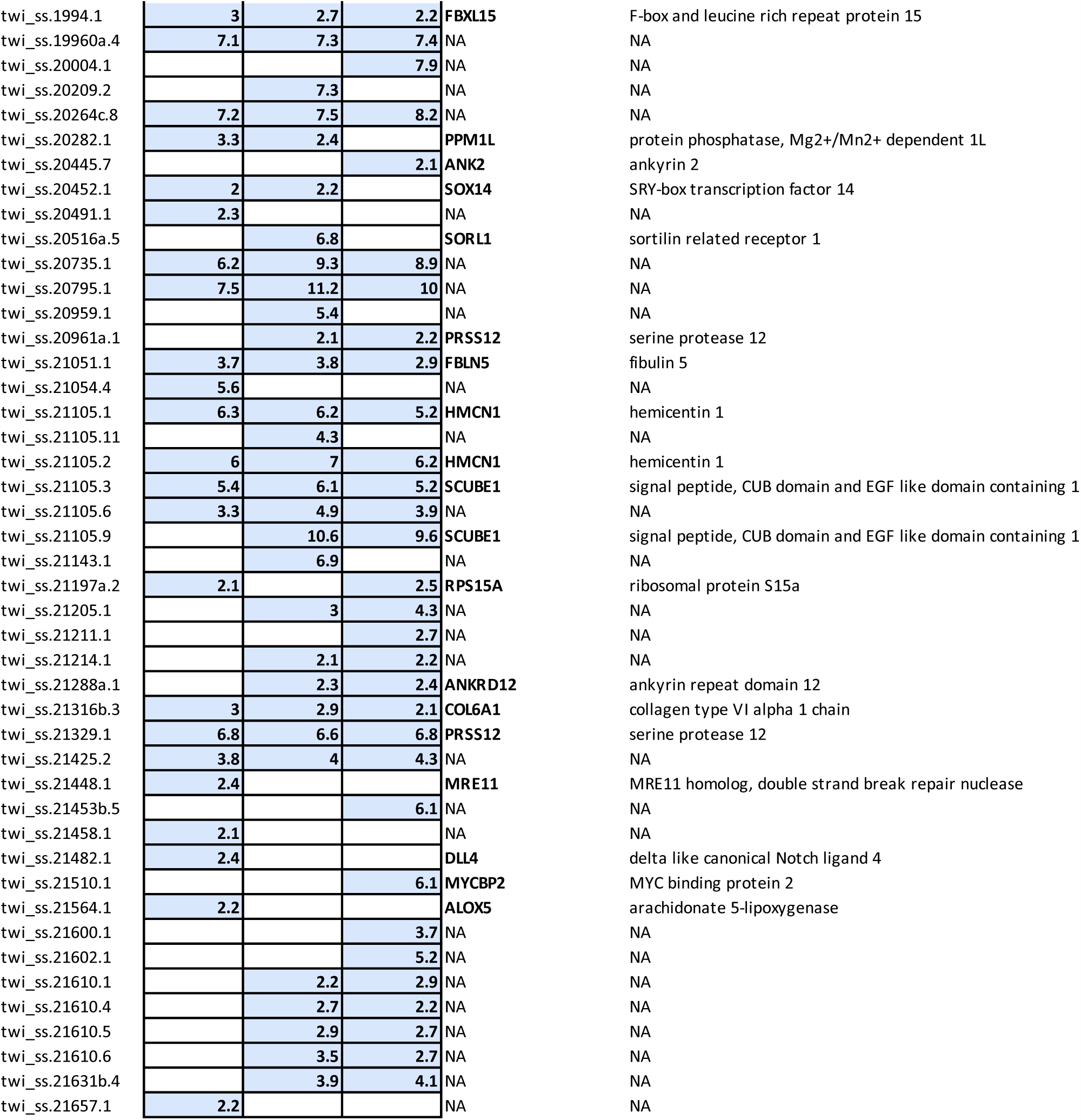
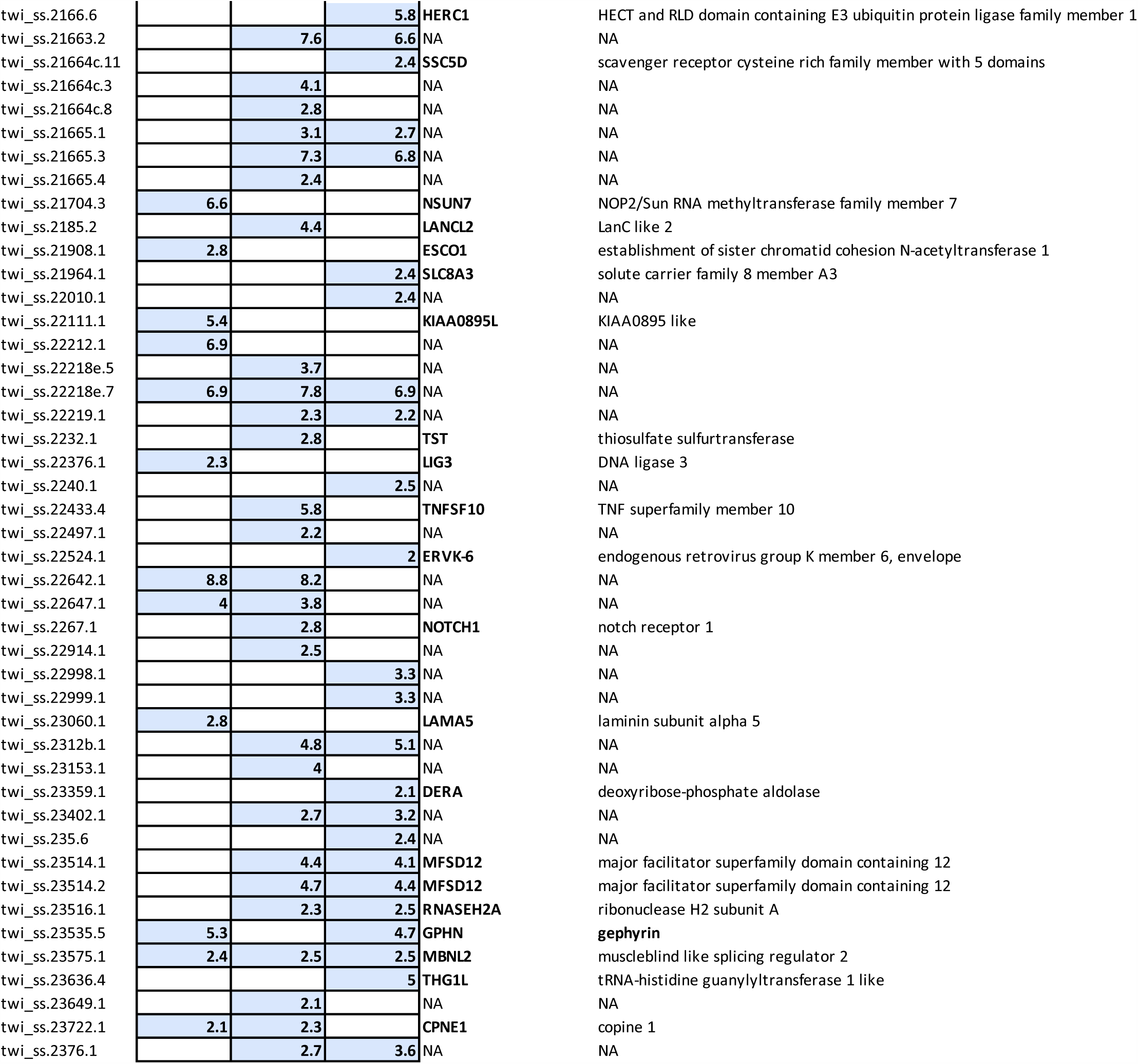
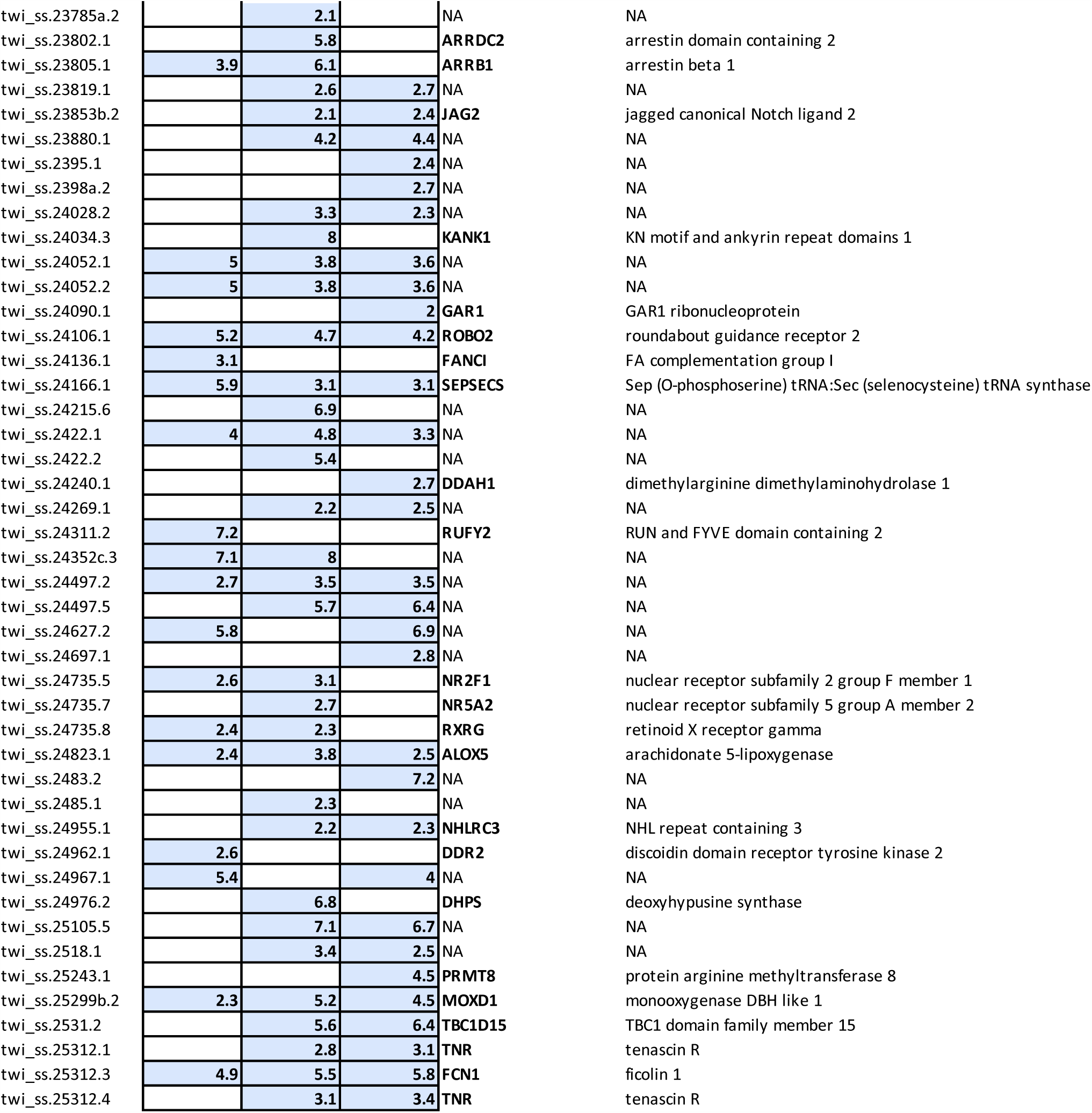
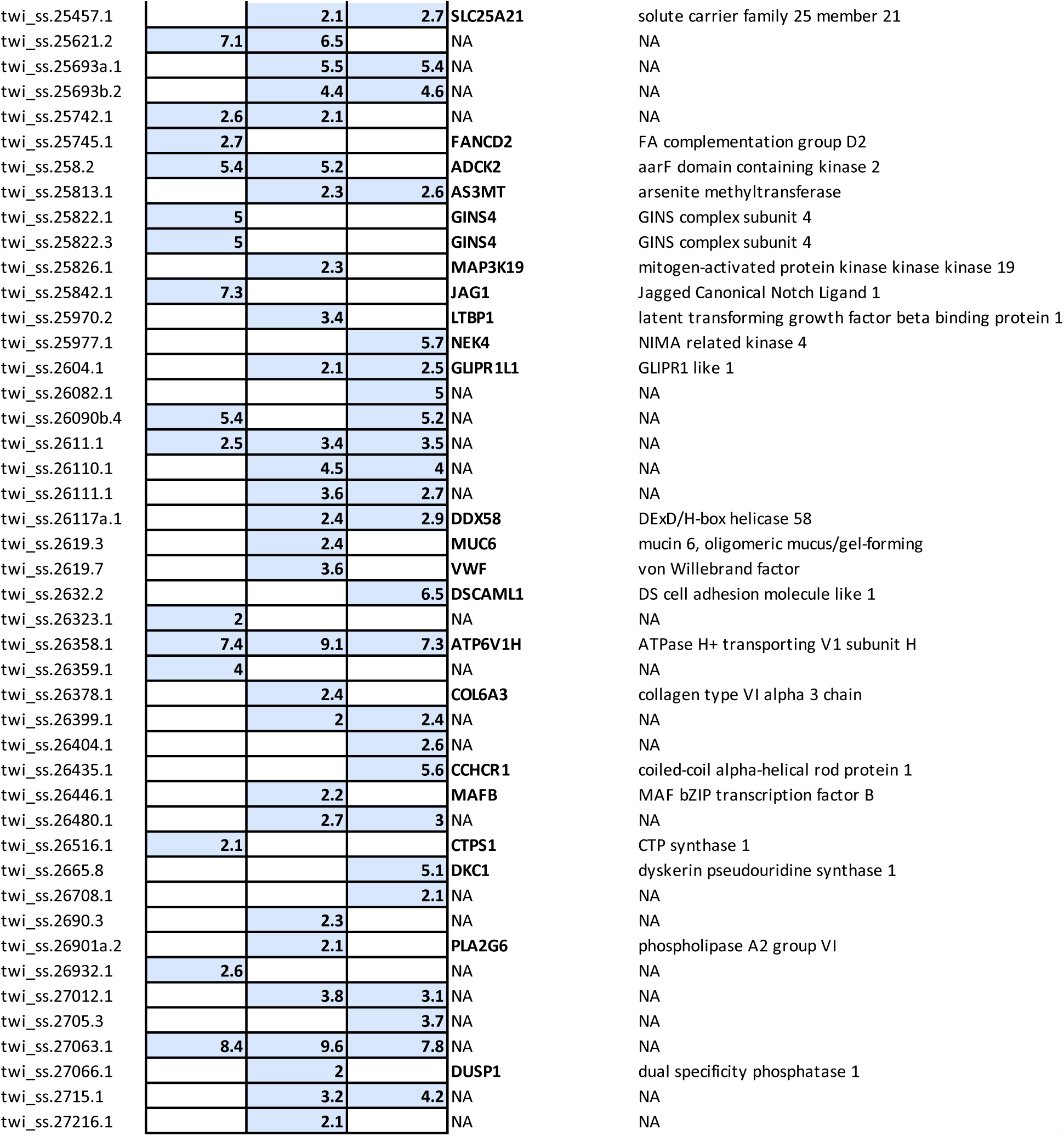
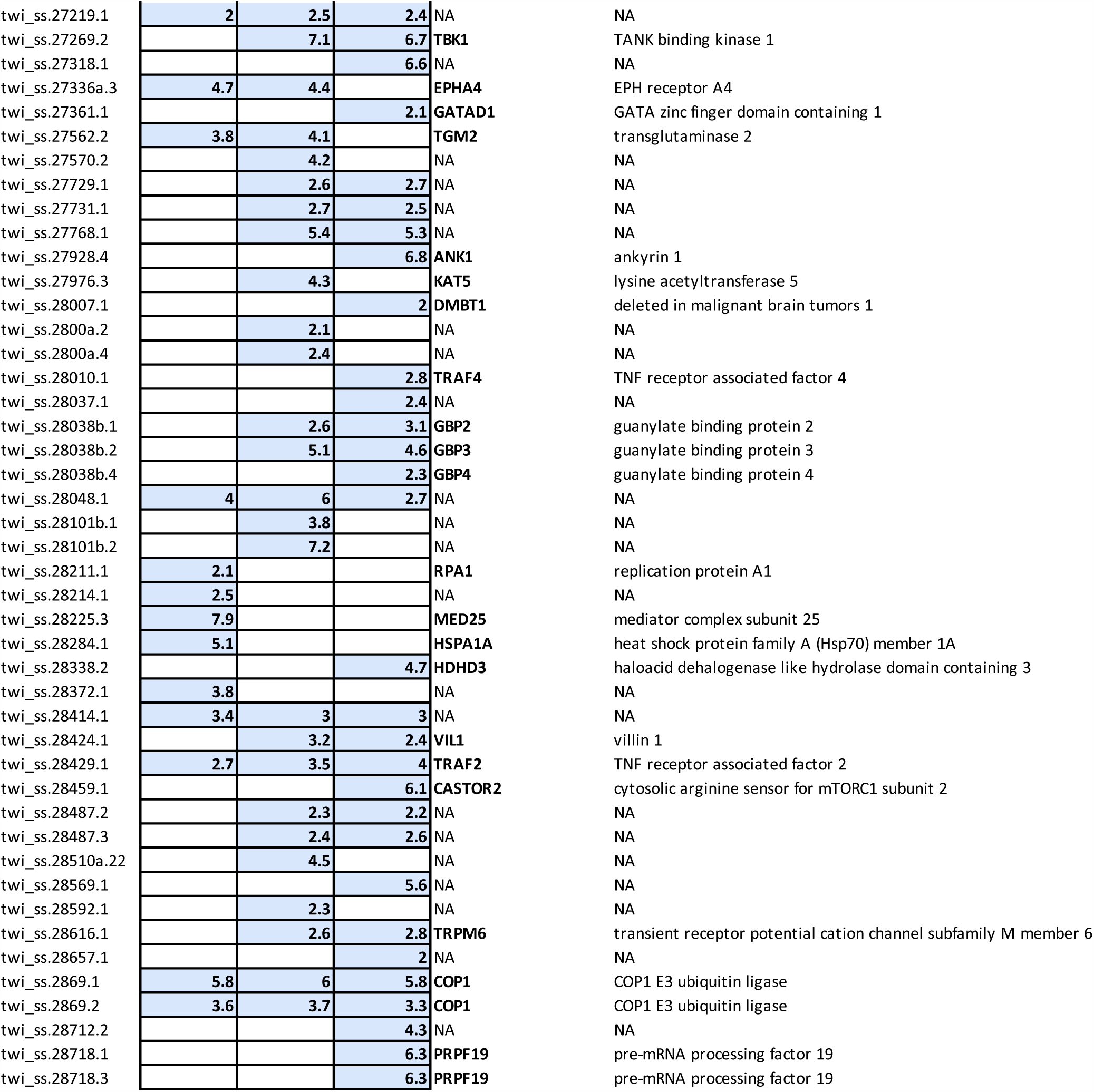
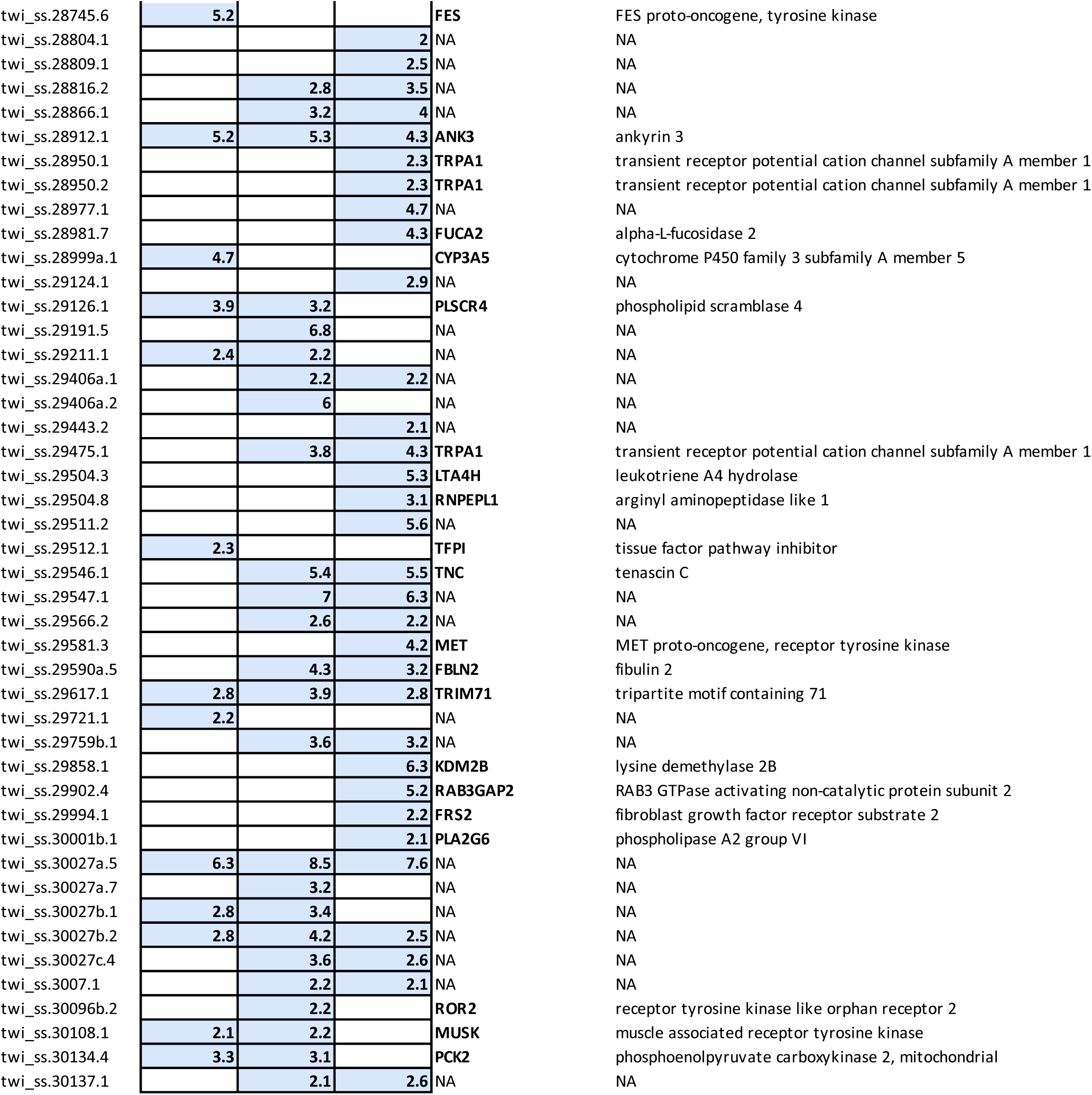
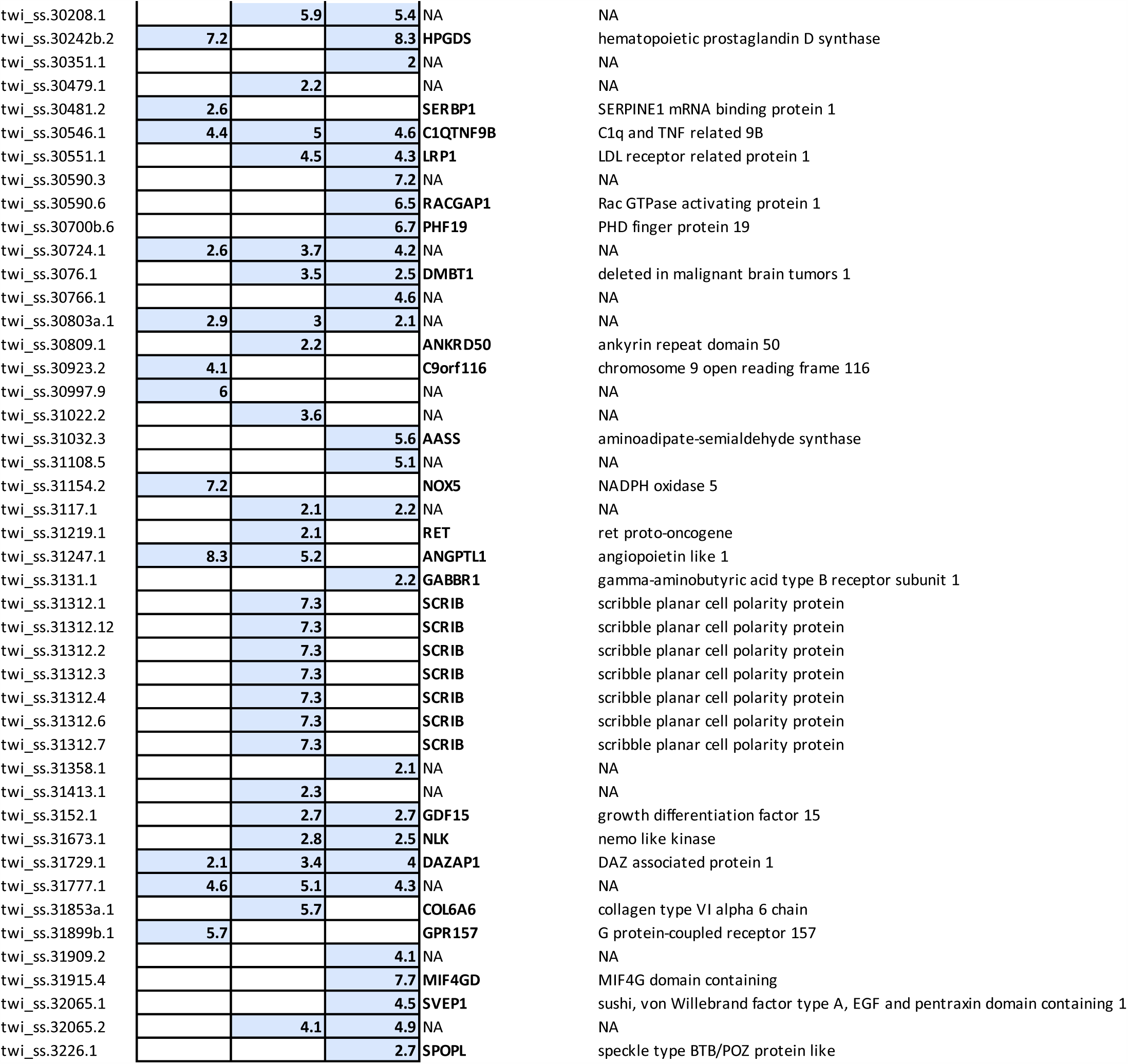
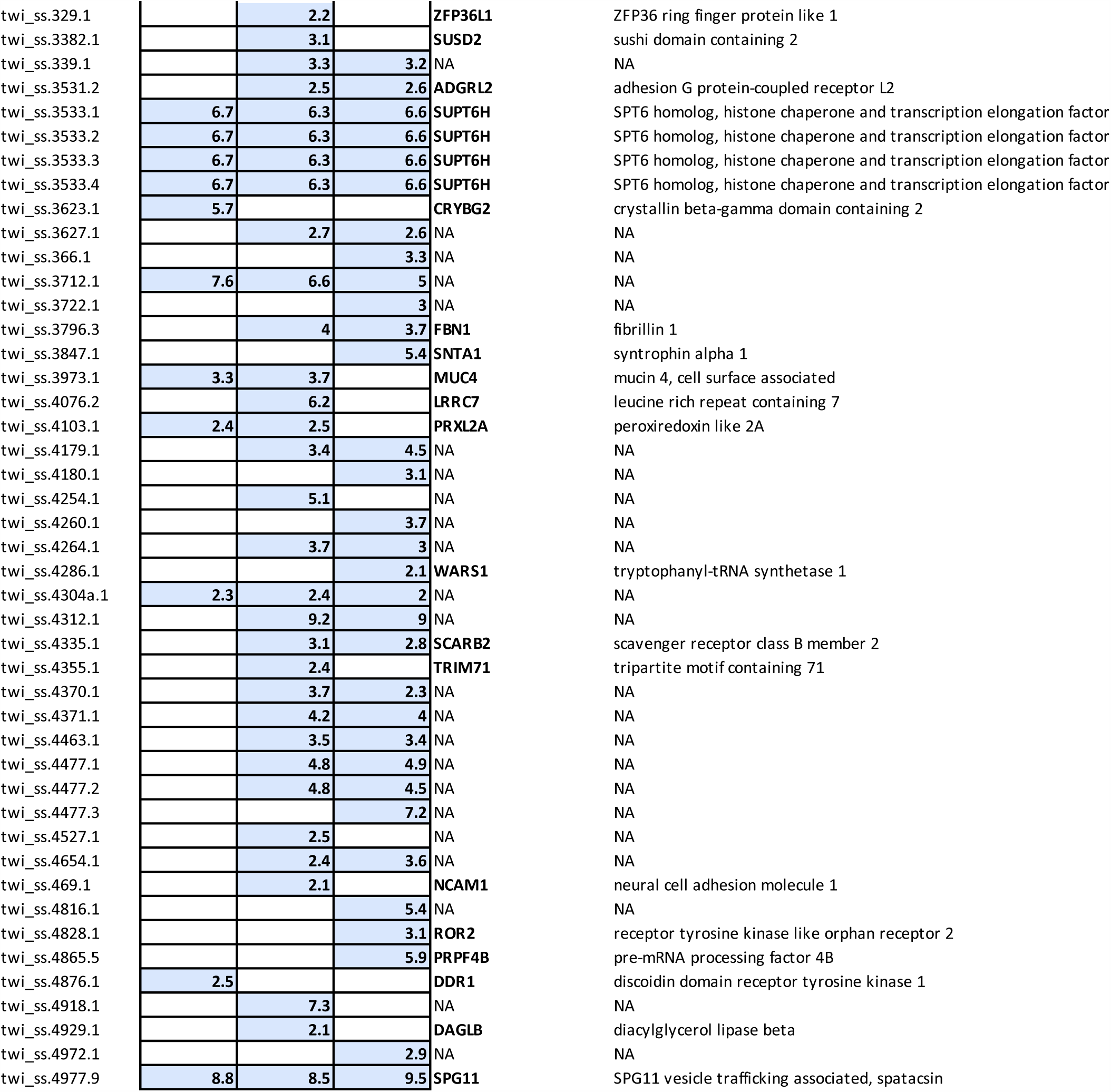
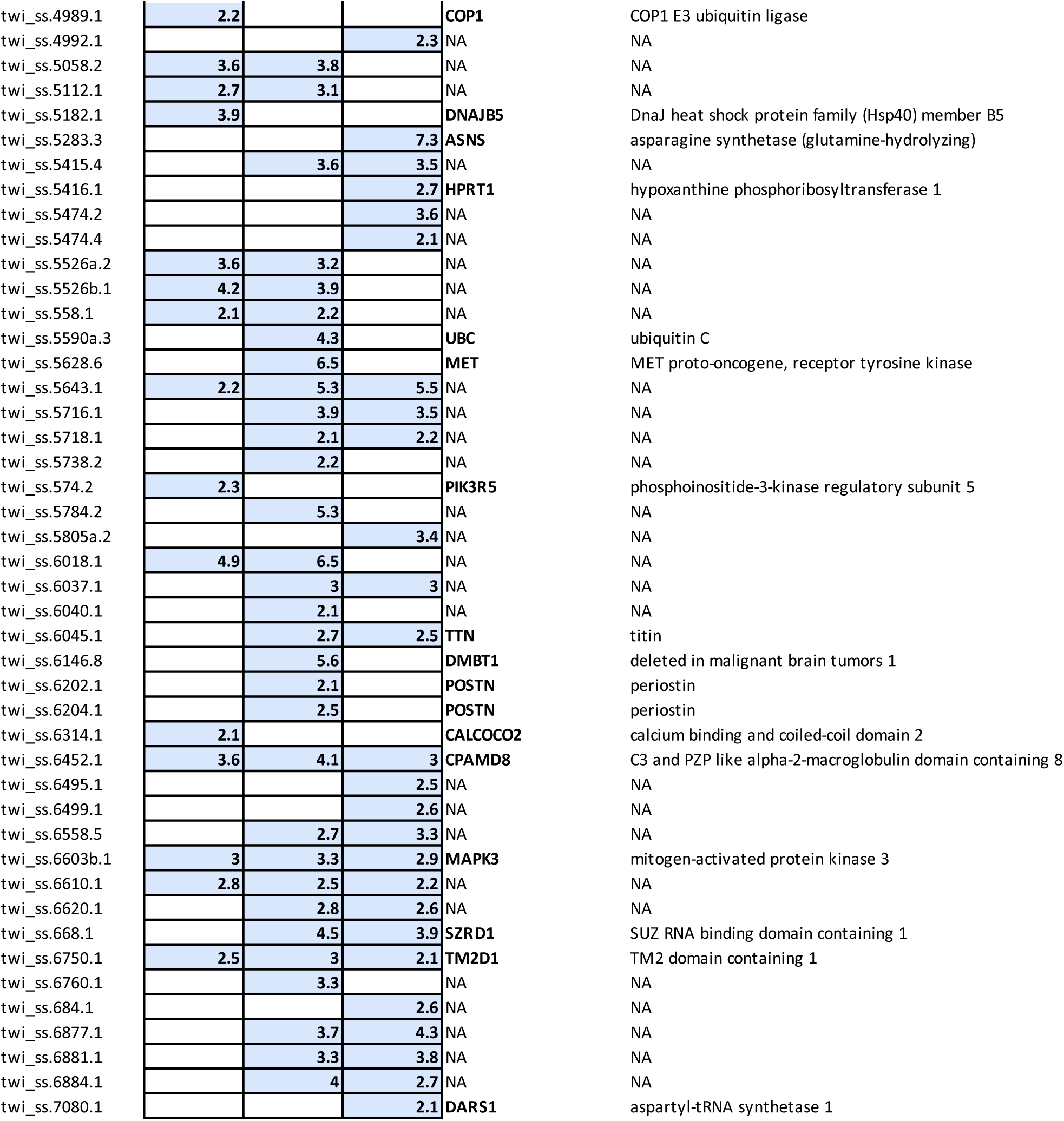
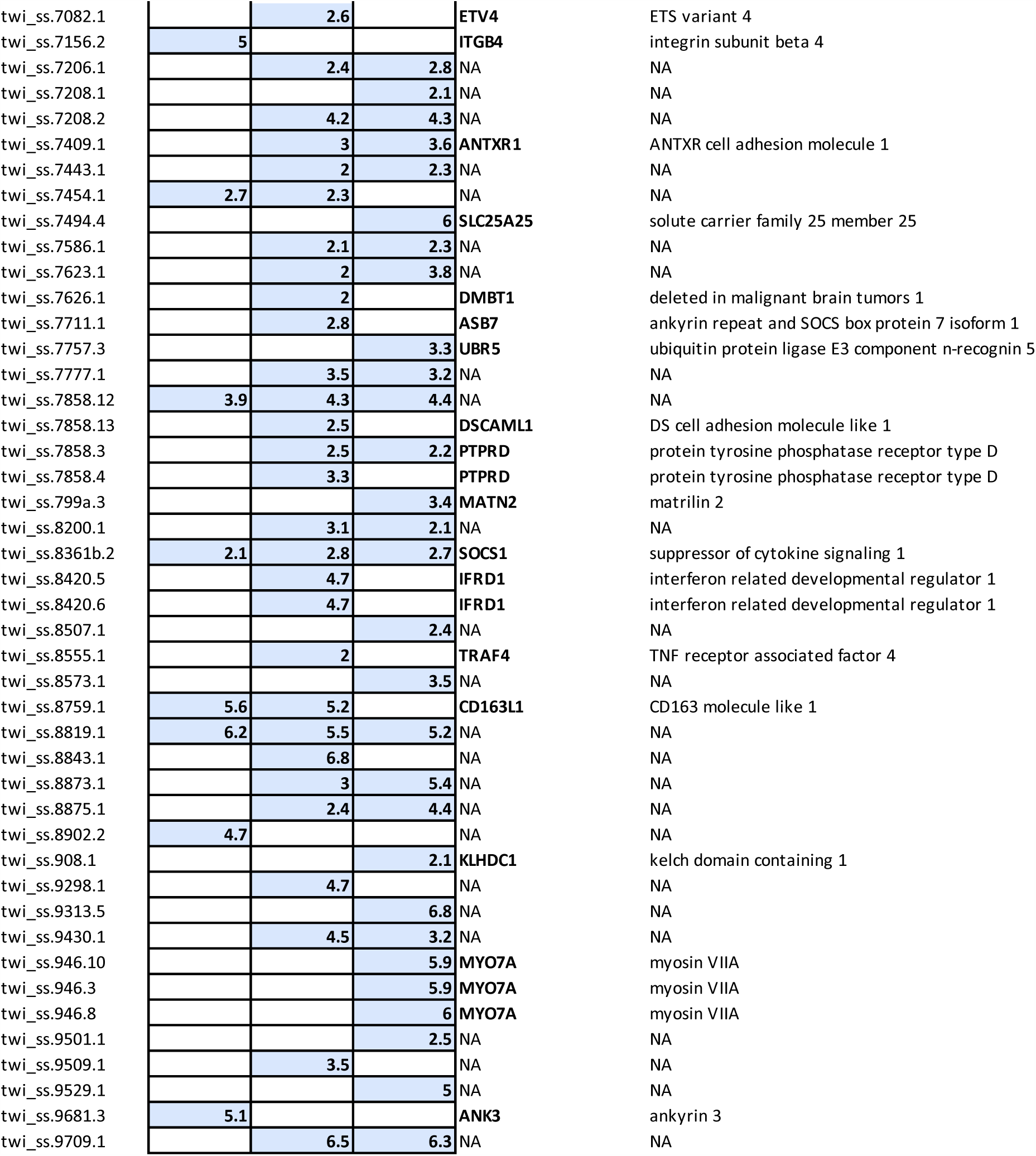
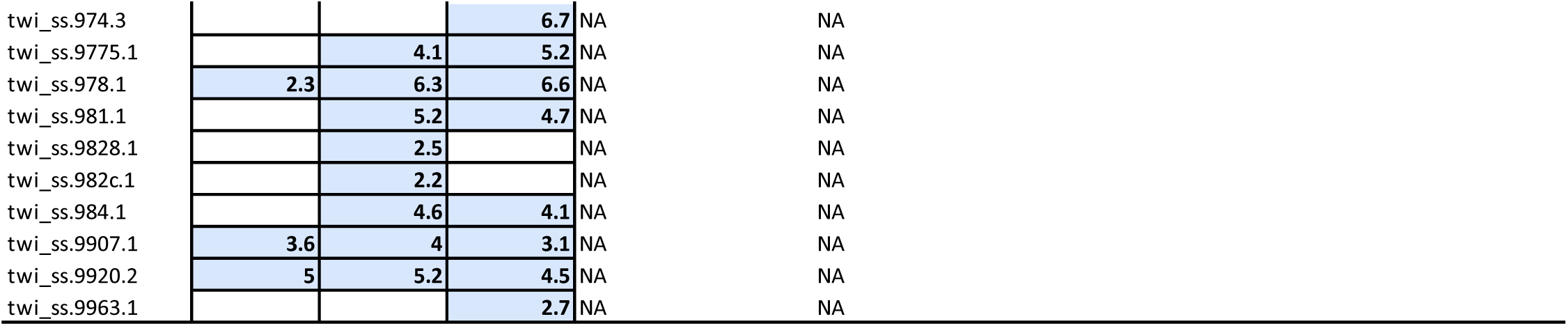
List of genes overexpressed 24 hours, 7 and 21 days after X-ray exposure. The table reports at least 2 log2-fold differentially expressed and statistically significant genes after multiple comparison correction (FDR<0.05) and their human homolog genes.

